# Cleavage of histone H2A during embryonic stem cell differentiation destabilizes nucleosomes to counteract gene activation

**DOI:** 10.1101/2021.08.10.455684

**Authors:** Mariel Coradin, Joseph Cesare, Yemin Lan, Zhexin Zhu, Peder J. Lund, Simone Sidoli, Yekaterina Perez, Congcong Lu, Elizabeth G. Porter, Charles W. M. Robert, Benjamin A. Garcia

**Affiliations:** Biochemistry and Molecular Biophysics Graduate Group, University of Pennsylvania, Philadelphia, PA 19104, USA; Department of Biochemistry and Biophysics, University of Pennsylvania, Philadelphia, PA 19104, USA; Epigenetics Institute, Perelman School of Medicine, University of Pennsylvania, Philadelphia, PA 19104, USA; Department of Biochemistry, Albert Einstein College of Medicine, Bronx, NY 10461, USA; Department of Oncology, St. Jude Children’s Research Hospital, Memphis, TN, USA; Department of Biochemistry and Molecular Biophysics Washington University in St. Louis, St. Louis MO 63110 USA; Department of Molecular, cellular and Developmental Biology, University of Colorado Boulder, Boulder CO 80309 USA

**Keywords:** Proteolysis, histone, stem cells, mass spectrometry, ChIP-Seq, differentiation, chromatin remodelers

## Abstract

Histone proteolysis is a poorly understood phenomenon in which the N-terminal tails of histones are irreversibly cleaved by intracellular proteases. During development, histone post-translational modifications are known to orchestrate gene expression patterns that ultimately drive cell fate decisions. Therefore, deciphering the mechanisms of histone proteolysis is necessary to enhance the understanding of cellular differentiation. Here we show that H2A is cleaved by the lysosomal protease Cathepsin L during ESCs differentiation. Using quantitative mass spectrometry (MS), we identified L23 to be the primary cleavage site that gives rise to the clipped form of H2A (cH2A), which reaches a maximum level of ~1% of total H2A after four days of differentiation. Using ChIP-seq, we found that preventing proteolysis leads to an increase in acetylated H2A at promoter regions in differentiated ES cells. We also identified novel readers of different acetylated forms of H2A in pluripotent ES cells, such as members of the PBAF remodeling complex. Finally, we showed that H2A proteolysis abolishes this recognition. Altogether, our data suggests that proteolysis serves as an efficient mechanism to silence pluripotency genes and destabilize the nucleosome core particle.

## Introduction

Eukaryotic cells condense their DNA around histone proteins to form chromatin. The basic repeating unit of chromatin is the nucleosome, consisting of approximately 150 base pairs (bp) of DNA wrapped around a histone octamer composed of two copies of each histone H3, H4, H2A and H2B^1^. Extending out from the nucleosome core, histone tails (residues 1-30) are the major acceptor of post-translational modifications (PTMs), among which acetylation, methylation and phosphorylation are the most common. Different biological outcomes are associated with the specific sites and types of histone PTMs, such as gene activation for H3K27ac, gene silencing for H3K9me3, and mitosis for H3S10ph^2^. The proper regulation of gene expression by histone PTMs is crucial for guiding cellular differentiation and dictating cell fate.

Embryonic stem (ES) cell differentiation gives rise to the three germ layers, the endoderm, mesoderm, and ectoderm, which are critical for development^3^. This process of differentiation is tightly regulated by external cues as well as internal machinery, such as transcription factors and epigenetic modulators, which have been shown to remodel chromatin and alter gene expression during cell fate commitment^4^. During lineage commitment, cells experience major changes in chromatin arrangement, resulting in alteration of gene expression^5,6^. As cells differentiate, chromatin becomes less accessible and more compact with decreased physical elasticity^7^. These changes are driven in part by chromatin remodelers acting to reposition nucleosomes and/or promote histone variant exchange^8^. Differentiation also leads to changes in histone PTM patterns^6^. For instance, during the early stages of differentiation, ES cells show high levels of acetylation, a mark of active chromatin, while the heterochromatic H3K9me3 is barely detectable in pluripotent ES cells ^9^.

Histone proteolysis, another form of regulation, is a less explored process in which histones tails are enzymatically cleaved, thereby removing existing PTMs and precluding future PTMs until histone exchange occurs. Histone proteolysis has been observed in different biological contexts. H2A, for example, has been shown to undergo proteolysis in acute monocytic leukemia at residue V114^10^. Similarly, H2B has been found to be cleaved in hepatocytes ^11^. Histone clipping appears to be dictated not only by the primary amino acid sequence, but also by the PTMs on histone tails. Jumonji-C domain-containing proteins JMJD5 and JMJD7 have been shown to cleave methylated arginine in mouse embryonic fibroblasts (MEFs)^12^. Furthermore, during osteoclastogenesis, H3K18ac has been shown to modulate proteolysis of H3 by matrix metalloproteinase 9 (MMP-9)^13^.

In the context of cellular development, the lysosomal protease Cathepsin L (CTSL) has been shown to cleave H3 during ESC differentiation^14^. CTSL localizes to the nucleus upon Ras activation, where it is known to cleave the CDP/Cux transcription factor, regulating the cell cycle ^15^. Additionally, CTSL has been shown to promote cellular senescence in IMR90 fibroblasts and primary melanocytes by cleaving the histone variant H3.3^16^. CTSL knockout embryos exhibit abnormal visceral endoderm formation^17^. Furthermore, CTSL-deficient mice show irregular hair follicle development and other skin pathologies^18^. Prior to our work, the biological significance of histone proteolysis by CTSL in regulating gene expression during stem cell differentiation had remained unresolved, as questions of whether this protease cleaves other histones and whether this processing affects nucleosome stability had yet to be investigated.

Here, we report that CTSL cleaves histone H2A during embryonic stem cell differentiation. We hypothesized that H2A proteolysis serves as a quick mechanism to remove acetylation on the N-terminal histone tail, thus counteracting gene activation during development. Using high-resolution quantitative mass spectrometry (qMS), we localized the primary cleavage sites to lie between amino acids 22-25. Additionally, we found that CTSL knockdown leads to a global increase in H2A acetylation after four days of differentiation. Using chromatin immunoprecipitation followed by next generation sequencing (ChIP-seq), we found that acetylated H2A is re-distributed to promoter regions upon CTSL knockdown. We have also discovered that members of the SWI/SNF remodeling complex bind to acetylated H2A but not cleaved H2A (cH2A). Lastly, we found that nucleosomes and dimers containing cH2A are less stable than full length H2A (FL-H2A), as revealed by protein degradation analysis. Taken together, our results uncover cellular consequences of histone proteolysis and describe the role of this cleavage in altering H2A modifications and nucleosome stability during cell fate commitment.

## Results

### The N-terminal tail of histone H2A is cleaved during mouse embryonic stem cell differentiation

Prior studies have reported that histone H3 is cleaved upon cellular differentiation^14^. To determine if differentiation involves the cleavage of other histones, we treated mouse embryonic stem cells (mESCs) with retinoic acid (RA) to induce their differentiation into embryoid bodies (EBs, Figure 1A). Using immunoblot analysis, we identified that histone H2A is cleaved upon differentiation at day one and four (Figure 1B), and no change was observed for H2B. Next, we sought to interrogate whether cleaved H2A (cH2A) still associates with chromatin by isolating mono-nucleosomes from both undifferentiated mESCs and EBs using micrococcal nuclease (MNase) digestion. As shown in Figure 1C, cH2A is only present in mono-nucleosomes derived from EBs, but not in undifferentiated cells (consistent with our results in Figure. 1B), suggesting that cH2A is chromatin associated.

**Figure 1.**
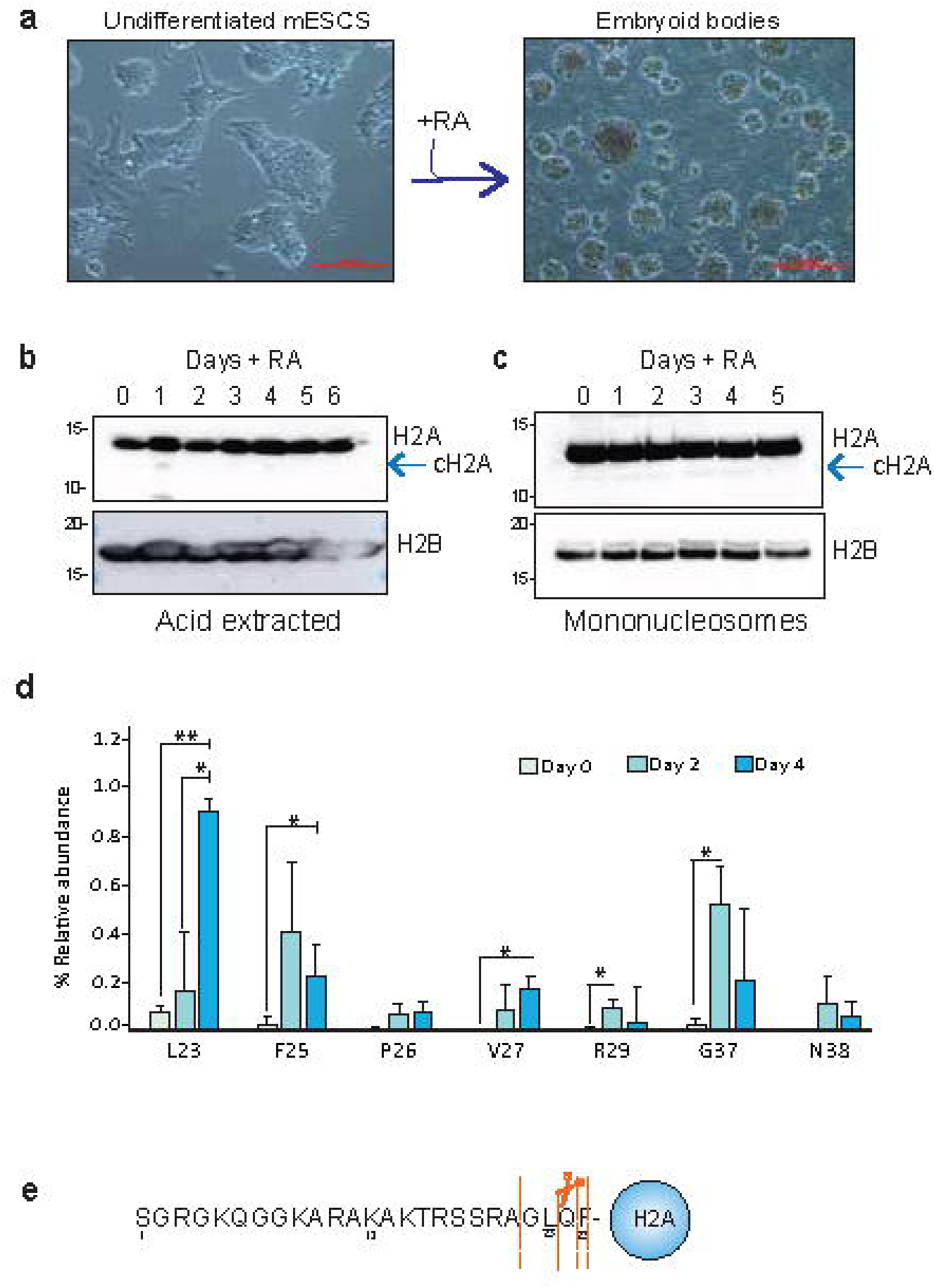
Histone H2A is cleaved during ES cell differentiation. **A** Undifferentiated ES cells, (left panel) from embryoid bodies (right panel) after treatment with Retinoic acid (RA) treatment. **B** Western blot analysis on acid extracted histone or **C** mononucleosomes during 6 or 5 days of differentiation, arrow indicates the presence of cleaved H2A (cH2A). **D** Mass Spectrometry quantification of cH2A relative abundance compared to FL-H2A. Asterisks indicate significant differences between undifferentiated (day 0) and differentiated cells (day 2 and 4), two-tailed student t-test (*p < 0.05, **p < 0.01). Bar plots represent the average of 3 biological replicates and the error bars represent the S.D **E** Illustration of the N terminal tail of H2A, dotted lines indicate secondary cleavage sites while solid line denotes primary and most abundant cleavage site.

Next, we utilized quantitative mass spectrometry to identify the cleavage site(s) on H2A and quantify the levels of cH2A upon differentiation. Histones from differentiated and undifferentiated mESCs were extracted with acid and then further purified by reverse phase liquid chromatography (RP-HPLC). Characteristic fractions corresponding to canonical H2A.1^19^ (Supplementary Fig. 1A) were analyzed by liquid chromatography coupled to MS (LC-MS). In agreement with our immunoblot data, we detected cleavage of H2A at day two of differentiation, which significantly increased further by day four of RA treatment. The most abundant cleavage site we found was at L23 with ~1% of all H2A being cleaved at this site (Figure 1, D and E). Other cleavage sites that increased through differentiation were also identified at F25, V27 and G37. Although additional cleavage sites were detected along the H2A tail (Supplementary Fig 1B-D), their abundance did not change during differentiation, indicating that these sites were not regulated during cell development.

Numerous variants of H2A exist, including macroH2A, H2A.Z, and H2A.X, which have different roles in cellular differentiation^20^. Although addressing whether H2A variants undergo cleavage during development is beyond the scope of this work, sequence alignment shows that the cleavage motif is conserved across the majority of the variants (Supplementary Fig 1E). It remains to be determined if these variants are being cleaved during differentiation or other biological processes.

### Cathepsin L facilitates H2A proteolysis upon cellular differentiation

Previously, we reported that CTSL cleaved H2A *in vitro* ^21^ and other studies have similarly found CTSL-mediated cleavage of histones *in vitro* using reconstituted nucleosomes^22^. Analysis of the cleavage sites of CTSL substrates has revealed a motif consisting of glutamine at the P1 position and aromatic residues at the P2 position^23^. The H3 cleavage motif Q/LAT (/ indicates cleavage site)^14^, is similar to where H2A is being cleaved (G/LQF) as both contain hydrophobic and aromatic amino acids (Figure 2A). Together, this led us to the hypothesis that CTSL serves as the H2A protease.

**Figure 2.**
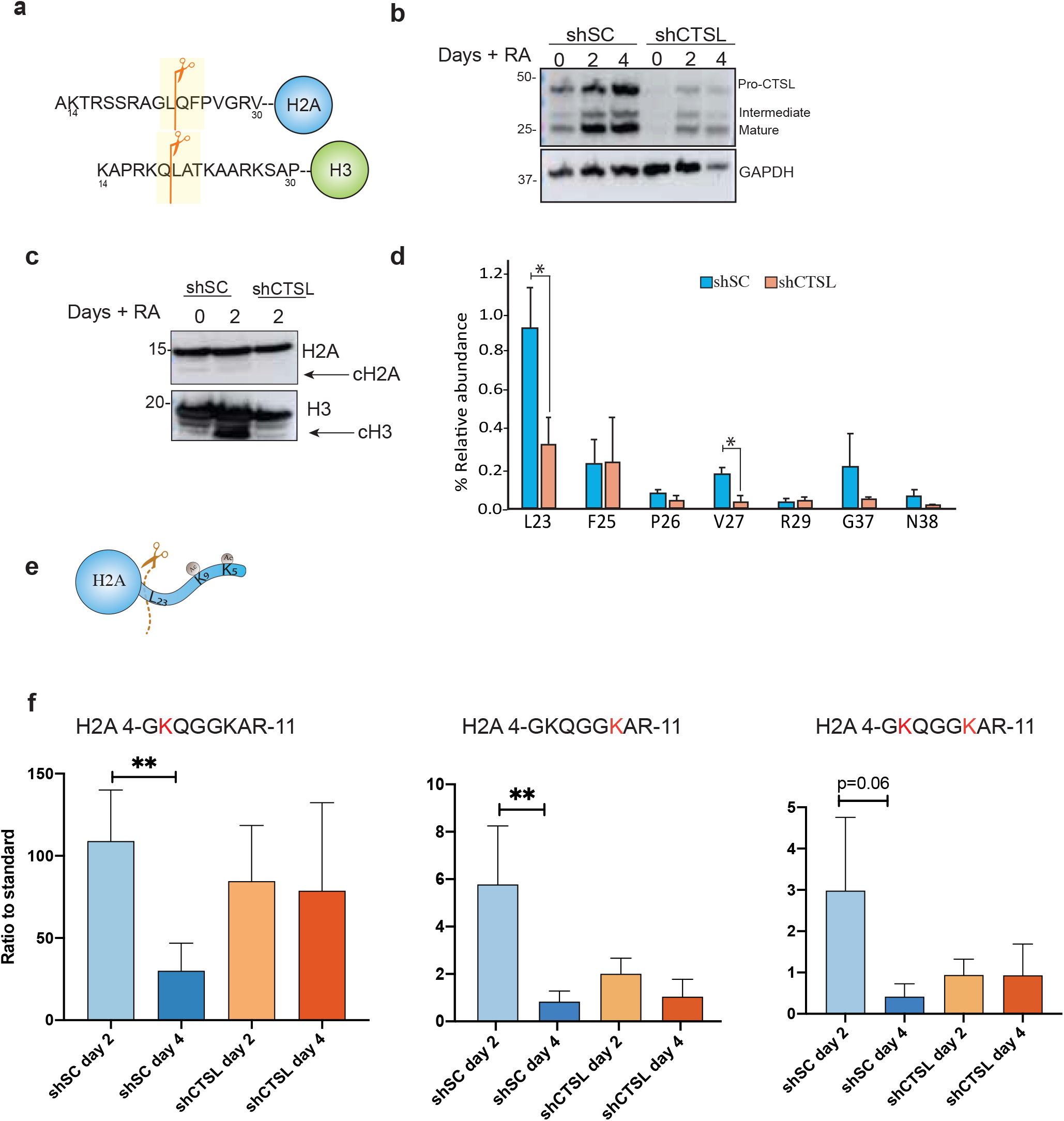
Cathepsin L facilitates H2A proteolysis upon differentiation. **A** Sequence similarity between H3 and H2A cleavage sites. **B** Validation of CTSL knockdown in undifferentiated and differentiated cells, GAPDH was used as loading control. **C** Western blots on acid extracted histones from undifferentiated WT mESCs and differentiated embryoid bodies with and without shCTSL **D** Mass spectrometry quantification of cH2A after 4 days of RA treatment. Asterisks indicate significant differences between scramble shRNA (shSC) or CTSL KD cells (shCTSL), two-tail student t-test (*p < 0.05) and at least 3 biological replicates in each time point. **E** Illustration of the post translational modifications on the N-terminal tail of histone H2A. **F** MS quantification of histone acetylation marks on H2A on differentiated ES cells. two-tailed student t-test (*p < 0.05, **p < 0.01) and N=3. All results represent the average of three biological replicates and the error bars represent the S.D

To investigate this further, we knocked down the expression of CTSL using shRNA (shCTSL) in mESCs (shSC was used as control) (Figure 2B). Next, we differentiated mESCs (shCTSL and shSC) for two days, acid-extracted histones and analyzed them by immunoblot. Although shCTSL mESCs still differentiated into embryoid bodies, cleavage of H2A was reduced compared to the control (Figure 2C). Similarly, when we blotted for total H3, we also found less cleavage of histone H3 (cH3) in shCTSL cells compared to control cells as well (Figure 2C). To quantify the levels of cH2A upon CTSL knockdown, we employed top-down MS to monitor intact H2A as well as the cleavage products. Again, we observed a significant reduction of cH2A when the expression of CTSL was reduced (Figure 2D and Supplementary Figure 2A and B). Proteolysis in the control cells was observed not only at L23 but also at other residues, including V27, suggesting that CTSL cleaves H2A at multiple sites. Nevertheless, the abundance of L23 was three-fold higher than V27, indicating that L23 is most likely the primary cleavage site for CTSL *in vivo* (Figure 2D).

To understand how cleavage of H2A by CTSL affects post-translational modification (PTM) patterns, we analyzed acetylation levels at H2AK5 and H2AK9, as these H2A N-terminal marks are associated with gene activation^24^. Our mass spectrometry data showed that the acetylation levels of H2AK5 and H2AK9 change during normal ESC differentiation. In undifferentiated cells, the levels of all possible forms of acetylated H2A (K5ac, K9ac and dually acetylated K5acK9ac) are approximately 10% (Supplementary Figure 2C-E). Two days after RA treatment acetylation levels increase significantly to 19%. Finally, after four days of RA treatment, acetylation levels decrease to 7%. To interrogate how CTSL proteolysis affects the levels of acetylated H2A during differentiation (Figure 2E), we quantified the levels of acetylated H2A in control and shCTSL cells after RA treatment by spiking in synthetic peptides corresponding to H2AK5ac, H2AK9ac, H2AK5acK9ac and the unmodified H2A peptide. As shown in Figure 2F, the levels of H2AK5ac, significantly decreased after four days of RA treatment in control but not in shCTSL cells. In fact, the levels of H2AK5ac were relatively unchanged. Similarly, H2AK9ac levels also did not decrease in shCTSL cells. Although at day two, the levels of H2AK9ac and the H2AK5acK9ac seemed lower in mESC shCTSL cells versus control cells at day two, the trend of acetylation levels not changing from day two to day four in shCTSL cells. Together, our data indicates that CTSL-mediated proteolysis serves to rapidly remove multiple acetyl marks on the H2A N-terminus, thereby potentially regulating gene expression during differentiation.

### Knockdown CTSL leads to genome wide redistribution of acetylated H2A in stem cells

Given that histone tails, specifically those of H3 and H4, have been shown to be dynamically modified during stem cell differentiation ^25^, and our observations indicating that H2A is also dynamically acetylated during differentiation, we asked how proteolysis regulated the genome-wide localization of H2A acetylation. We performed ChIP-seq for H2AK9ac in shSC and shCTSL mESCs as well as in differentiated embryoid bodies (Figure 3A). Alterations in the total levels and genomic distribution of H2AK9ac in undifferentiated CTSL KD cells compared to control cells were not apparent by MS or ChIP-seq (Supplementary Figure 3A-B). In agreement with our MS data, we found a genome-wide decrease in H2AK9ac on day two of treatment when CTSL was knocked down but not at day four. (Figure 3B). Moreover, when determining the genomic distribution of H2AK9ac, we found a 4% increase at promoters in CTSL KD EBs (Supplementary Figure 3D). Next, we performed differential binding analysis, here we found that ~1.1% of peaks changed significantly between day two and day four in the control (Log_2_ fold change >1 (up) or < −1 (down), FDR < 0.05), which was not observed upon knockdown of CTSL (Figure 3D). This led to the question of whether the expression of those genes are altered upon CTSL knockdown. To answer this question, we performed RNA-seq in shSC and shCTSL cells.

**Figure 3.**
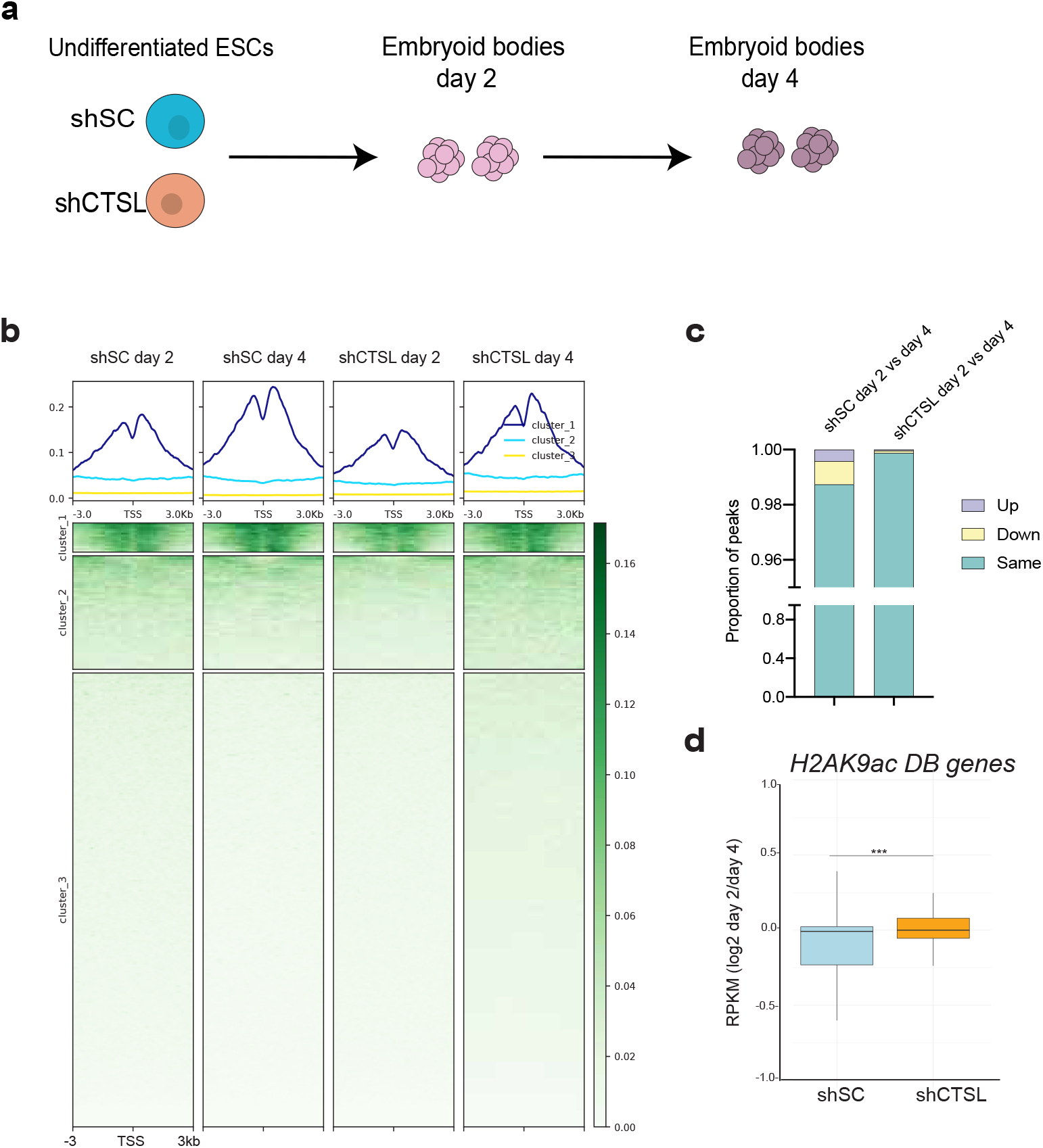
Genome wide localization of acetylated H2A **A** Schematic of differentiation time points used for ChIP-seq and RNA-seq analysis. **B** Heatmap showing enrichment of H2AK9ac over mm10 genes centered around the TSS at day 2 and day 4 of EBs formation in control cells (shSC) or CTSL KD cells (shCTSL). **C** DiffBind^56^ analysis comparing H2AK9ac occupancy at day 2 vs day 4 in shSC and shCTSL **D** Gene expression by RNA-seq. The boxplot indicates the log2 fold change in gene expression from day 2 vs day 4 in shSC (blue) and shCTSL (orange) using genes (n=347)with significantly lower H2AK9ac occupancy (by DiffBind). Wilcoxon-rank sum two-sided was used to determined significance (p=0.0001).

We hypothesized that genes that lose H2AK9ac after four days of RA treatment will also have lower gene expression compared to day two. As demonstrated in Figure 3D, genes (n=347) with significantly less H2AK9ac have lower gene expression after four days of differentiation in the control but not upon CTSL knockdown. Interestingly, gene ontology (GO) analysis showed that genes with significantly decreased H2AK9ac in WT cells (n=347) at day four of RA treatment are involved in cellular differentiation and nervous system development (Supplementary Figure 3C). Taken together, these data suggest that H2A proteolysis by CTSL aids in gene regulation by silencing genes involved in pluripotency while activating genes to promote cell linage commitment.

### Acetylated H2A is recognized by PBAF complex in mESCs

Changes in the H2A acetylation patterns upon loss of CTSL expression likely alters the recruitment of regulatory proteins to chromatin. In addition to H2AK9ac, our MS data showed increased levels of H2AK5ac in differentiated cells upon CTSL KD (Figure 2F and Supplementary Figure 2C). This mark is associated with gene activation^24,26^, but its reader protein remain to be identified. Thus, to identify potential readers, we used synthetic peptides corresponding to H2AK5ac, H2AK9ac and H2AK5acK9ac as bait in peptide pulldown assays. Isolated proteins were characterized by LC-MS/MS with further validation by immunoblot (Figure 4A). As a negative control, we used an H2A peptide with only N-terminal acetylation since H2A is known to be co-translationally acetylated at the N-terminal serine residue by NatD^27,28^. All other H2A peptides also include this N-terminal acetylation.

**Figure 4.**
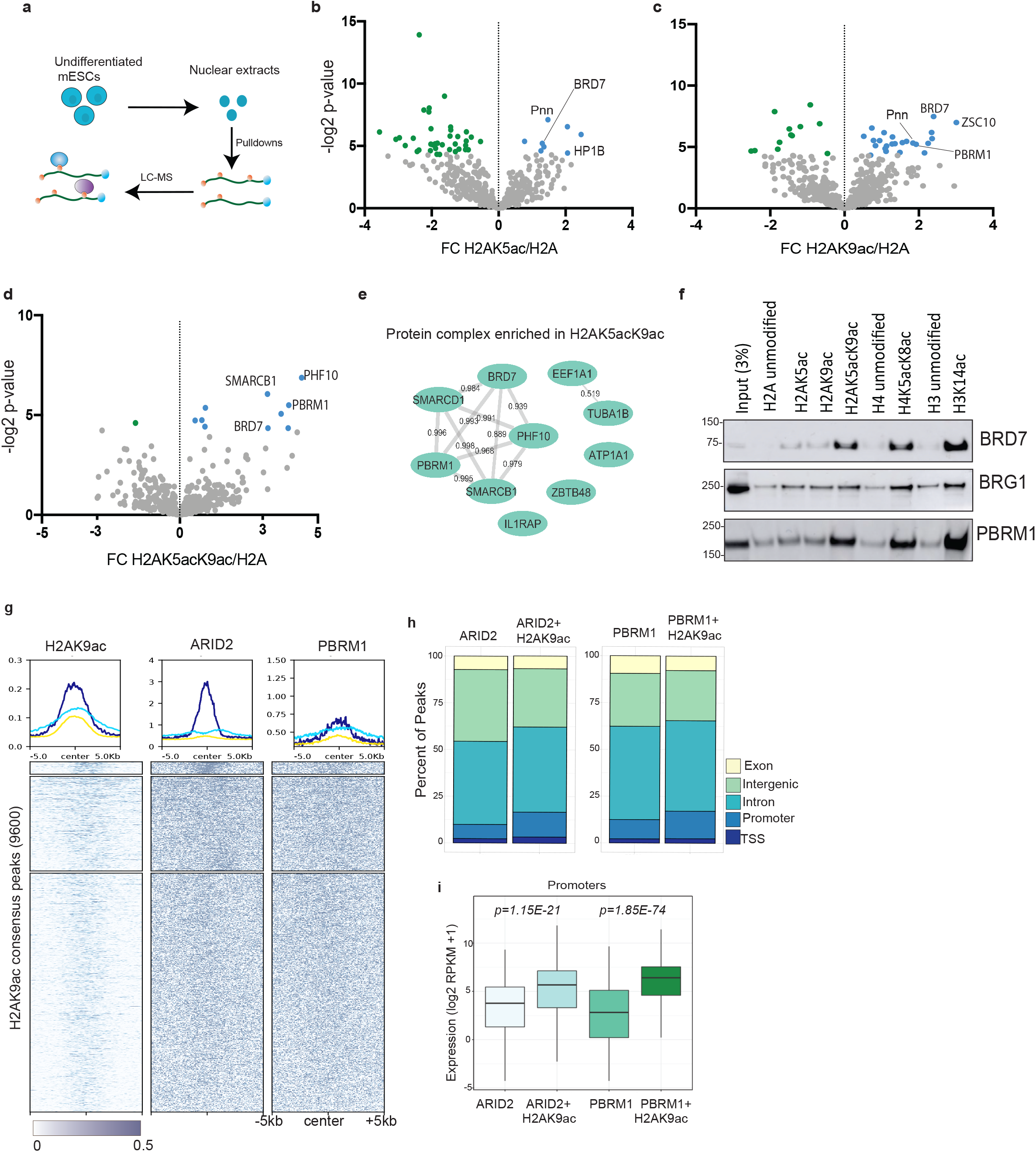
Members of pBAF recognized acetylated H2A. **A** Representation of histone peptide pulldown experiments using nuclear extracts. Volcano plots showing the fold change between unmodified H2A (control) peptide and (**B)** H2AK5ac, (**C**) H2AK9ac or (**D**) H2AK5acK9ac. Proteins significantly enriched by acetylated peptides versus the control peptide are labeled in blue (Student t-test). **E** STRING analysis showing associations between the proteins enriched in H2AK5acK9ac pulldown, the numbers represent the interaction strength. **F** Western blots validation of MS results. **G** Heatmap showing enrichment of Arid2 and Pbrm1 on H2AK9ac consensus peaks in pluripotency. **H** Genome wide distribution of pBaf and pBAF/ H2AK9ac colocalized regions. **I** Expression levels (log2 RPKM+1) of promoter in pBaf and pBaf/H2AK9ac dual regions, Significance was determined using two-sided unpaired student t-test.

The H2AK5ac and H2AK9ac peptide baits succeeded in capturing members of the Polybromo-1 BRG1 Associating Factor (PBAF) chromatin remodeler complex, including Brd7 and Pbrm1 (Figure 4B, C). Similarly, the dually acetylated H2AK5acK9ac was also able to pull down PBAF members, such as Smarcd1, Phf10, Pbrm1 and Brd7. Enrichment of PBAF proteins was more prominent when H2A was acetylated at K5 and K9 simultaneously (Figure 4D, Supplementary Figure 4A). Similarly, Brd7 preferentially binds to dually acetylated H2A (Figure 4F). However, it appears recombinant Brd7 binds better to H2AK5ac alone *in vitro* (Supplementary Figure 4B), suggesting that additional proteins may enhance recognition of dually acetylated H2A. Furthermore, we found that Brg1 showed similar affinity for all acetylated peptides (Figure 4D and Supplementary Figure 4A). Since histone H4 is also known to be acetylated at its N-terminal tail, which resembles the sequence of H2A, we reasoned that acetylated H4 may also interact with the PBAF complex. Repeating our affinity pulldown experiments using unmodified and a dually acetylated (K5acK8ac) H4 peptides, we observed enrichment of PBAF compared to an unmodified control (Figure 4F and Supplementary Figure 4B). However, as expected, the main reader for H4K5acK8ac, Brd4 ^25,29^ attained greater enrichment than Brg1 and Brd7 (Supplementary Figure 4B-C). Pbrm1 has been previously reported to bind H3K14ac^30^, which we likewise observed in our experiments. However, we found that Pbrm1 also binds to H2AK5acK9ac (Figure 4F). Additionally, we observed that TBP binds acetylated H2A and H4, highlighting its positive correlation with gene activation (Supplementary Figure 4D).

We next performed ChIP-Seq for PBAF specific proteins (Pbrm1 and Arid2) to establish genes that are co-occupied by H2AK9ac and PBAF in mESCs. Our data showed that 54% of all Arid2 peaks contain H2AK9ac(n=9257). Similarly, 55% of all Pbrm1 peaks are also marked with H2AK9ac (n=8316) (Figure 4G and Supplementary Figure 4E and F). When we compared the PBAF occupied regions without H2AK9ac to where they coexisted, we found a remarkable increase of PBAF at gene promoters with acetylated H2A present (Figure 4H). For instance, 7.6% of peaks exclusive to Arid2 are found at the promoter, however when acetylated H2A is present, we observed 13.3% of peaks correspond to promoter regions. Likewise, we noticed a 4% increase of Pbrm1 localization at promoters in presence of acetylated H2A, suggesting that H2AK9ac facilitates PBAF localization to promoters. To correlate H2AK9ac and PBAF cooccupancy, we compared the expression of genes containing H2AK9ac and PBAF at their promoters to those exclusively found for PBAF. As shown in Figure 4I, expression of genes that are co-occupied by PBAF and acetylated H2A have significantly higher expression than those only marked by PBAF (P-value<0.0001), suggesting that potential recruitment of PBAF by H2AK9ac promotes gene expression.

### H2A proteolysis prevents PBAF recognition of acetylated H2A

Since the modifications on histone tails can regulate the recruitment of proteins to chromatin, proteolytic severing of the tails could preclude these interactions normally mediated by modifications on the tails. We hypothesized that removal of the H2A tail, and therefore H2A tail acetylation, would disrupt binding of PBAF to H2A. To test this, we performed coimmunoprecipitation followed by MS (IP-MS) analysis using mESCs expressing either FLAG-tagged full-length H2A (FL-H2A) or N-terminally truncated H2A, representing cleavage at L23 (cH2A). Inspection of the interactome of FL-H2A compared to cH2A showed 96% of the proteins interacted with both cH2A and FL H2A (Figure 5A). However, in agreement with our previous findings, members of the PBAF chromatin remodeler complex (Smarcd1 and Actl6a) were only found to interact with FL-H2A and not cH2A. Additionally, we found that Importin-9 (Ipo9) was enriched in cH2A sample compared to FL-H2A (Figure 5B and C). Ipo9 is known to translocate H2A-H2B dimers from the cytosol to the cell nucleus ^31^. We also found that NPL1 (nucleosome assembly protein 1, also known as Nap1), which is known to exchange H2A/H2B dimers ^32^, was enriched by cH2A over FL-H2A. Interestingly, we found Cbx1, 3 and 5 to preferentially interact with FL-H2A (Figure 5B and C). Gene ontology analysis showed that the proteins enriched by cH2A are typically associated with RNA splicing and gene expression while those enriched with FL-H2A are involved in nucleosome assembly and DNA packaging (Supplementary Fig. 5A-B).

**Figure 5.**
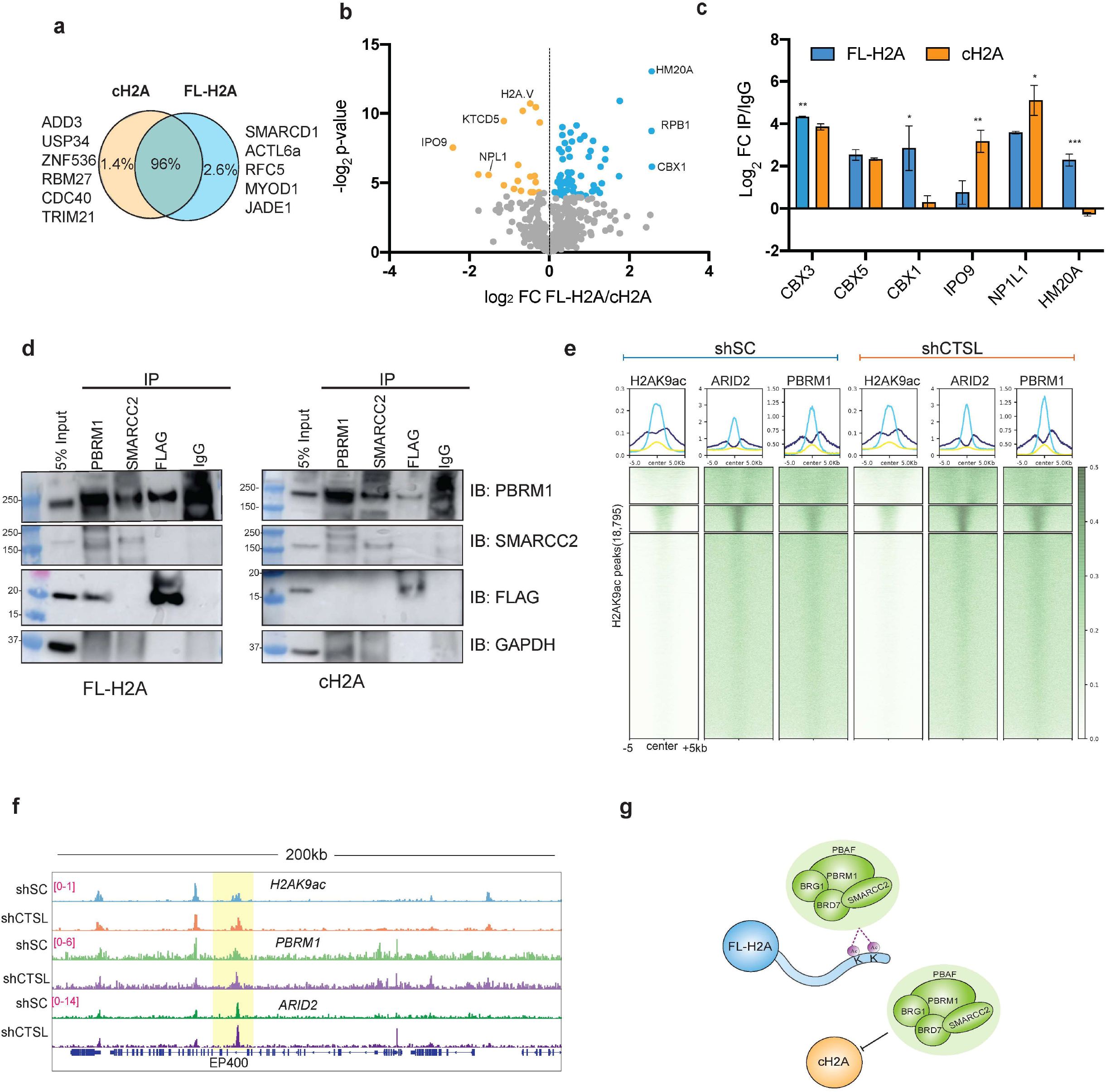
H2A proteolysis prevents PBAF recognition of acetylated H2A **A** Ven diagrams showing overlapping and unique proteins identified by IP-MS. **B**. Volcano plots showing the fold change between FL-H2A (blue) and cH2A (orange) Significantly enriched proteins by Student t-test are highlighted in blue or orange. Only proteins significantly enriched over IgG (FC >2 and p-val >0.05) were used for analysis **C** Bar plot of proteins differentially enriched in each condition, showing the average of three biological replicates Student t-test; *p < 0.05, **p < 0.01, ***p < 0.001. The error bars represent the SD **D** Co-IPs using whole cell extracts of mESCs expressing either FLAG-tagged full-length H2A (FL-H2A) or FLAG-tagged cH2A (cH2A). **E** Heatmap showing enrichment of Arid2 and Pbrm1 on H2AK9ac peaks in shSC and shCTLS after four days of RA treatment **F** Genome browser snapshot of H2AK9ac, Arid and Pbrm1 over EP400. **G** Recognition of acetylated H2A by the PBAF before and after histone proteolysis

We next validated our IP-MS results by doing co-immunoprecipitation assays followed by immunoblotting. As shown in Figure 5D and Supplementary Figure 5C, FL-H2A coprecipitated with Pbrm1 more so than cH2A. Additionally, Pbrm1 but not Smarcc2 (Baf170) coprecipitated with FL-H2A, suggesting that Pbrm1 interacts with H2A acetylation through its bromodomains, which bind to acetylated lysine residues. Given that proteolysis removes the binding substrate for PBAF, we hypothesized that Cathepsin L KD cells should have higher PBAF occupancy at H2AK9ac sites. To test this hypothesis, we focused on day four as cH2A abundance is at its maximum. We performed ChIP-Seq for Pbrm1 and Arid2 in shSC and shCTSL cells at day four of RA treatment. When we analyzed the occupancy of PBAF at H2K9ac peaks, we noticed an increased occupancy of both Arid2 and Pbrm1, (Figure 5E-F), indicating that PBAF remains bound to acetylated H2A when CTSL levels are reduced. Taken together, these results indicate that the PBAF protein complex recognizes acetylated H2A and that H2A proteolysis abrogates this recognition (Figure 5G).

### cH2A is associated with marks of active transcription and fast turnover

Our IP-MS results found a reduced interaction between CBX proteins and cH2A. Cbx1, also known as heterochromatin protein 1 β, plays an important role in gene silencing through its interaction with methylated H3K9^33^. This suggests that nucleosomes enriched with cH2A are depleted of histone H3K9me marks. To examine this possibility, we purified mononucleosomes containing cH2A and analyzed their PTMs by MS (Supplementary Figure 6A). As expected, nucleosomes with cH2A were mostly depleted of mono-, di- and tri-methylation at H3K9 compared to those with FL-H2A (Figure 6A, B). Instead, we found that cH2A-containing nucleosomes had higher levels of histone marks associated with gene expression, such as H3K14ac and H3K36me3 (Figure 6A). Taken together, this data suggests that FL-H2A can coexist with H3K9me in certain parts of the genome, while cH2A is most likely to be temporarily associated with accessible chromatin, due to its precursor, FL-H2A, being hyperacetylated on promoters of active genes.

**Figure 6.**
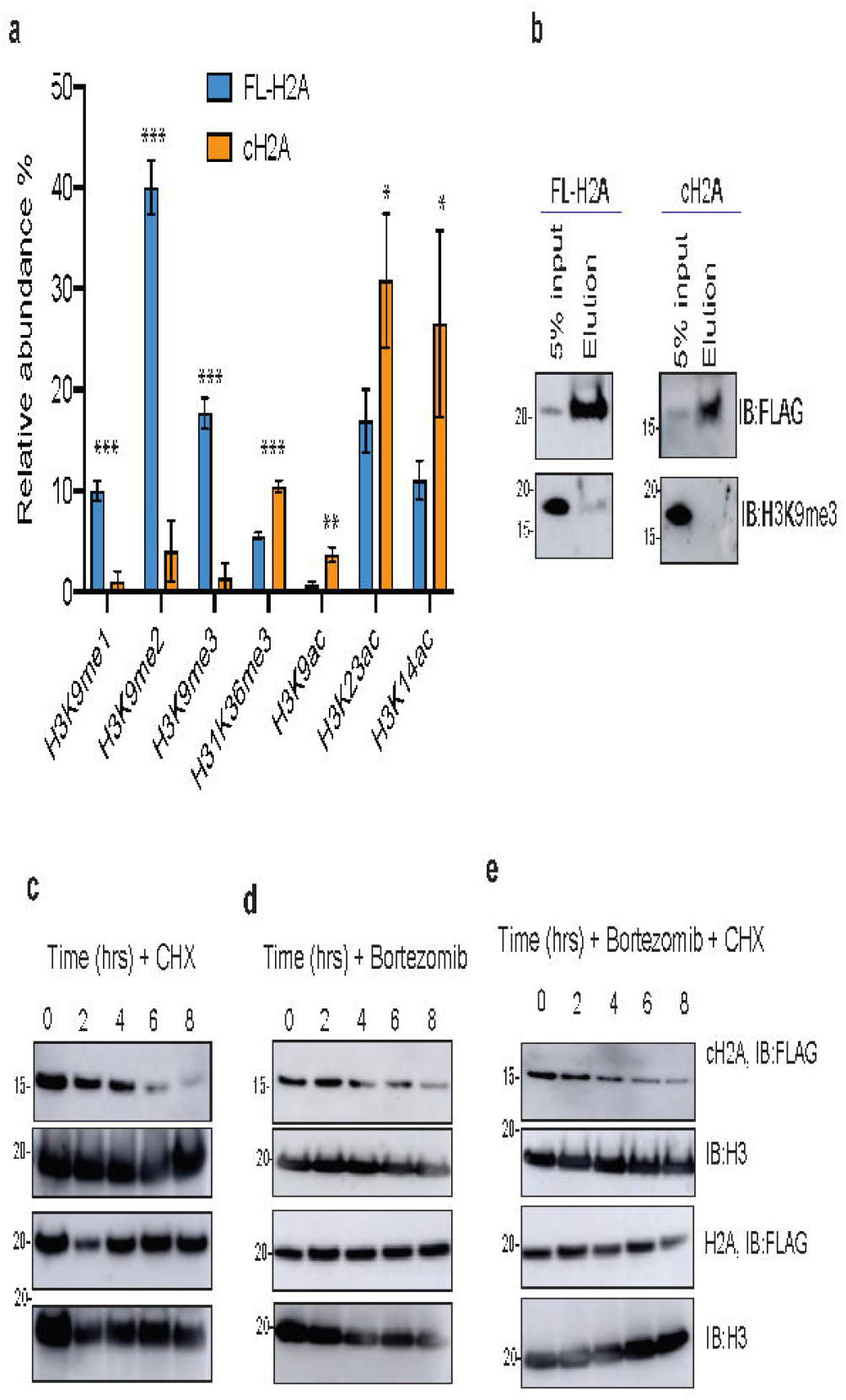
cH2A is associated with marks of active transcription and fast turnover. Mononucleosome IP-MS quantification of histone PTMs enriched in FL-H2A **A** or cH2A. significance is denoted by asterisk (*). Student t-test;*p < 0.05, **p < 0.01, ***p < 0.001. **B** MS validations by western blot. Protein stability assay using **C** cycloheximide 100ng/ml **D** or bortezomib (10nM) or both **D**. Western blots against FLAG and H3 in nuclear extracts. All experiments were done in biological triplicates

Aside from affecting histone modifications, cleavage of H2A by CTSL could also affect nucleosome structure and stability. Our recent publication using hydrogen deuterium exchange coupled with mass spectrometry (HDX-MS), showed that in a nucleosome context, the N-terminal tail of H2A is protected from exchange^34^, which is consistent with the finding that histone tail proteolysis destabilize the nucleosome *in vitro* ^35^. The loss of the N-terminal tail of H2A likely disrupts DNA-histone interactions that play important roles in the maintenance of nucleosome structure. Indeed, during cellular processes such as replication and transcription, the nucleosome undergoes conformational changes to allow access to DNA. Interestingly, we found that cH2A preferentially interacts with Npl1 (Figure 5B and C), a protein known to exchange H2A/H2B dimers. Thus, we hypothesized that cH2A-containing nucleosomes are readily exchanged or evicted compared to FL-H2A-containing nucleosomes. To measure the stability of cH2A in cells, we blocked protein synthesis with cycloheximide. We found that while FL-H2A protein levels remain stable for at least eight hours, cH2A undergoes rapid degradation (Figure 6C). Notably, in our IP-MS experiment, we found that cH2A but not FL-H2A interacted with Trim21 (Figure 5A), a ubiquitin ligase known to target proteins for proteasomal degradation ^36^. To investigate whether cH2A was being degraded by the proteasome, cells were treated with the proteasome inhibitor bortezomib and cH2A stability was subsequently monitored for eight hours. As shown in Figure 6D, the levels of cH2A still decrease after bortezomib treatment. This was further confirmed by treating cells simultaneously with cycloheximide and bortezomib (Figure 6E), indicating that cH2A may be degraded by another cellular degradation pathway. Overall, our data suggest that histone proteolysis occurs in open chromatin regions to remove key histone PTM sites and disrupt binding interactions, thus facilitating nucleosome destabilization and eviction, and promoting gene silencing of pluripotency related genes.

## Discussion

Histone proteins undergo a wide variety of changes to their PTM patterns to regulate gene expression. The addition of these chemical moieties, predominantly on lysine and arginine residues, serves to modulate electrostatic interaction between histones and DNA or recruit transcriptional regulators, such as chromatin remodeling complexes that can evict or slide nucleosomes along the DNA. Compared to classical histone PTMs, such as acetylation and methylation, the connection between histone proteolysis and gene expression in less clear. The first identification of histone clipping was reported three decades ago, involving cleavage of histone H1 *Tetrahymena* micronuclei^37^. Since then, histone proteolysis has been noted in additional organisms such as sea urchin^38^, chicken^39^, and yeast^40^. These studies contribute to the idea that histone proteolysis represents a mechanism of gene regulation similar to canonical histone PTMs. Indeed, Duarte et al. found that histone H3 proteolysis during cellular senescence promotes silencing of cell cycle related genes^16^.

In the context of cellular development, CTSL has been shown to cleave histone H3 ^14^, but the downstream effects of this cleavage event remain unclear. Complementing this study, we now report that histone H2A is also cleaved by CTSL during embryonic stem cell differentiation. Using high resolution mass spectrometry, we demonstrated that H2A undergoes proteolysis at amino acid L23, accounting for almost 1% of the entire H2A population in differentiated cells (Figure 1D). Although we found other cleavage sites in undifferentiated cells, they were not responsive to CSTL knockdown, which suggests the possibility of other H2A proteases beyond CTSL. In addition to cleavage at L23, we also found a secondary cleavage site at V27, which is significantly reduced upon CTSL knockdown (Figure 2D). This was not surprising given that on the histone H3 tail, CTSL is known to target a primary site (A21), however secondary sites exist.

One hypothesis is that CTSL first cleaves at L23, located right before the alpha 1-helix, and then proceeds to cleave again at V27 (within in the alpha one helix, further destabilizing the nucleosome by weakening DNA-histone interactions).

*In vitro* studies have found that CSTL is more active when H3 is acetylated at H3K18ac. However, dual acetylation at K18 and K23 has the opposite effect ^14^. Given that histone H2A is also acetylated within its N-terminal tail, we sought to interrogate how proteolysis affects H2A acetylation *in vivo.* We anticipated that changes in acetylation due to proteolysis would be difficult to monitor due to the stoichiometry being only 1%. Therefore, instead of using our standard histone PTM analysis platform ^41^, we designed a more sensitive method to monitor levels of H2A acetylation using synthetic internal standards and multiple reaction monitoring (MRM) MS, which revealed higher levels of acetylated H2A increased upon CTSL KD (Figure 2F).

It is important to reiterate that the overall abundance of H2AK5ac is roughly 8% in undifferentiated cells, which doubles two days after RA exposure (15%). By day four, the level of this mark decreases to ~9%. A similar trend is observed with H2AK9ac, which increases from 1% in undifferentiated cells to 2% in early differentiation before decreasing back to 1% in late differentiation. Since these observations are at the global level, localizing these changes more precisely in the genome may aid in uncovering how H2A acetylation regulates gene expression during cellular development. Furthermore, investigating how proteolysis affects genome-wide distribution of acetylated H2A will elucidate how this novel process contributes to cell fate decisions. We hypothesized that genes regulated by H2A proteolysis would retain H2A acetylation after four days of differentiation and concurrent knockdown of CSTL. We performed ChIP-Seq in control and CSTL KD mESCs. Surprisingly we found a redistribution in the genomic localization of H2AK9ac. After four days of RA treatment, the proportion of H2AK9ac peaks in promoter regions increased by 4% (Supplementary Figure 3D). Examining the expression of genes with significantly lower occupancy of H2AK9ac after four days of differentiation compared to day two, we found significantly higher levels of gene expression upon CTSL KD. This suggests that proteolysis promotes gene silencing as expected. In line with the findings from Duarte *et al.*^16^, we propose that proteolysis could serve as a rapid mechanism to silence genes involved in pluripotency, allowing cells to proceed through differentiation.

Our data indicates that H2A acetylation plays a key role in driving gene expression during development. Prior studies have found that H2A acetylation is associated with gene activation in *Drosophila* ^42^. Acetylation on H2AK5ac and H2AK9ac is known to be deposited by Tip60 (Kat5) ^43^. In ESCs, Tip60 is known to activate genes involved in proliferation and cell renewal ^44^. While the role of this acetyltransferase has been extensively described, little is known about H2A acetylation in ESCs and the readers of this mark. Our data showed that members of the PBAF SWI/SNF chromatin remodeler complex recognized different forms of acetylated H2A and had higher affinity for the dual acetyl mark, H2AK5acK9ac (Figure 4). As shown in Figure 4, PBAF members also seem to bind to H4K5acK8ac. This can be explained by the sequence homology between the N-terminal tails of H4 and H2A. Consistent with prior findings, we confirmed that BRD4 is the main reader of H4K5acK8ac and not H2AK5acK9ac. As previously reported, we also observed H3K14ac to interact with Brd7 and Pbrm1 ^30^. A possible explanation is that H3K14ac and acetylated H2A co-exist in the same nucleosome or in hyperacetylated chromatin regions. We also cannot exclude the possibility that Pbrm1 could accommodate all acetyl sites given its six tandem bromodomains. Despites this possibility, our ChIP-Seq experiments further support our initial observation of PBAF binding to acetylated H2A. As demonstrated in Figure 4G, we uncover a genome wide interaction between PBAF and H2AK9ac. Furthermore, we noticed an increased occupancy of PBAF at promoters that are occupied by H2AK9ac (Figure 4H), and these co-occupied genes have higher expression levels compared to those only occupied by PBAF. Together, our results suggest that H2AK9ac promotes gene expression by recruiting PBAF to promoter regions.

The PBAF complex has been shown to play a key role in maintaining pluripotency ^45^. In addition to H3K14ac ^46^, members of the BAF family are known to also interact with H3K4me1 ^47^. Given that PBAF also recognizes acetylated H2A, we propose that this interaction is diminished upon H2A proteolysis. Indeed, utilizing immuno-affinity precipitation followed by MS (IP-MS) and validation by immunoblotting, we found that FL-H2A but not cH2A is able to interact with the PBAF complex. We further supported this hypothesis by performing ChIP-Seq in shCTSL EBs, here we observed increased occupancy of PBAF members at H2AK9ac sites when we knockdown CTSL. Additionally, we also found that CBX proteins interact preferentially with FL-H2A over cH2A. Interactions between CBX proteins and FL-H2A could be mediated through the association of FL-H2A with H3K9 methylation. As demonstrated in Figure 6, FL-H2A-containing nucleosomes show enrichment for H3K9 methylation. Conversely, cH2A nucleosomes were found to be more associated with active marks, such as H3K9ac and H3K14ac, suggesting that histone proteolysis occurs in euchromatic regions. Finally, we also found that cH2A is less stable than FL FL-H2A (Figure 6), suggesting that proteolysis may destabilizes the nucleosomes *in vivo* and promote nucleosome eviction.

Taken together, our studies uncover a novel mechanism of gene regulation during cell fate determination. We propose a model (Figure 7) in which CTSL mediates the proteolysis of H2A to remove acetylation on the N-terminal tail, thereby silencing genes involved in the maintenance of pluripotency. This silencing is accomplished by preventing the recruitment of PBAF to acetylated H2A, which abrogates chromatin remodeling at these genes and blocks the binding of transcriptional machinery.

**Figure 7.**
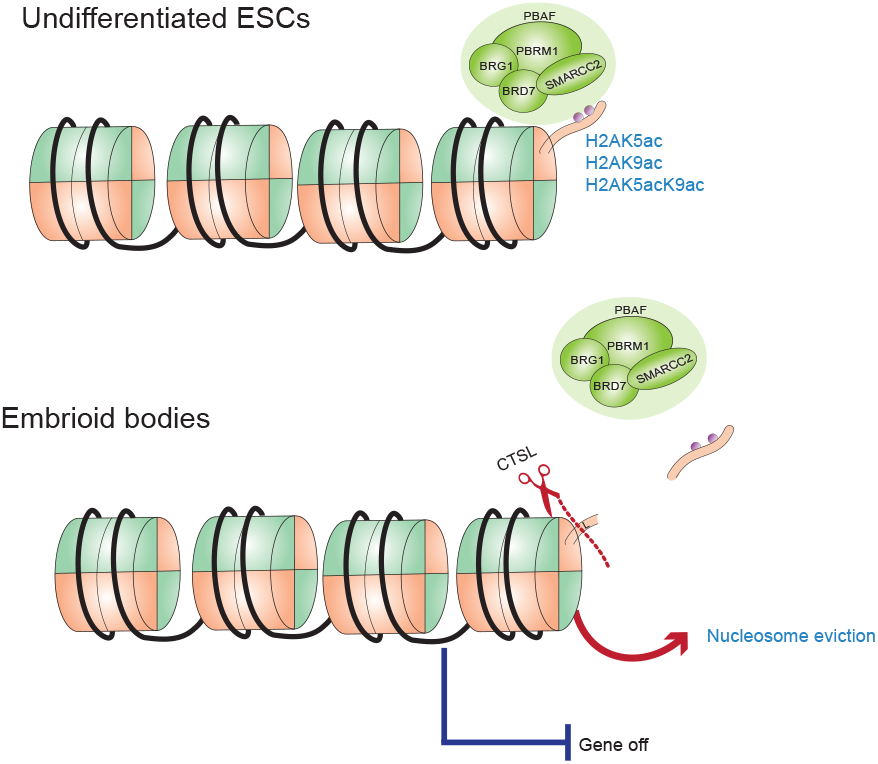
Proposed model of gene regulation by H2A proteolysis.

## Methods

### Cell Culture

CCE-Nanog-GFP MESCs were donated by Ihor Lemischka. Cells were grown at 37°C in 5% CO2 using DMEM with high glucose and sodium pyruvate (Thermo), supplemented with 15% characterized FBS (Hyclone,), non-essential amino acids (MEM) (Thermo), GlutaMax (Thermo), 10μM 2-mercaptoethanol (Thermo) and LIF(Millipore). Cells were grown on 0.1% gelatin (Millipore) coated plates and GFP + cells were sorted by flow cytometry before downstream experiments. For differentiation into embryoid bodies formation, ~1.5×10^7^ cells were plated in 10-cm dishes in media without LIF and supplemented with 10μM all-trans retinoic acid (Sigma). Cells were kept with constant rotation for 4 days and media was changed every other day. All cell lines were tested for mycoplasma.

#### Plasmids, cloning and generation of stable cell lines

To generate CTSL KD and Flag-tagged H2A (FL and cH2A) mESCs, approximately 4×10^5^ HEK293T cells were transfected with packaging vectors pPAX2 and VSGV (kind gifts from Dr. Shelley Berger) along with 1μg of FL-H2A or cH2A, or shRNA targeting CTSL. shRNA plasmid for CTSL KD (in pLKO.1) was obtained from the high throughput core at Penn and scramble control was a gift from Dr. Shelly Berger. To generate FLAG-Tagged FL-H2A and cH2A tagged with FLAG at the C-terminus, DNA fragments were cloned into pLenti plasmid (Origene) (EcoRI and PspXI). HEK293T were transfected in mESCs media (without LIF), and virus particles were collected for 3 days. Prior to transduction, the viral supernatant was passed through filtered using 0.45μm filter. Polybrene (Santa Cruz Biotechnology) was added to a final concentration of 8 ug/ml. Cells were infected for 24 hours followed by selection with 1μg/ml of puromycin (Thermo) for 5 days. Transduction efficiency of FL-H2A and cH2A was validated by immunoblotting with FLAG antibody (Sigma). CTSL knockdown efficiency was confirmed by RT-qPCR and immunoblots, with GAPDH as control in both cases. To assess protein stability, mESCs expressing FL-H2A and cH2A were treated with 100μg/mL cycloheximide (Sigma), 10 nM Bortezomib (Sigma)or a combination of both drugs for eight hours with timepoints collected every two hours. Cell pellets were snap frozen in liquid nitrogen and stored at −80°C.

### Histone isolation and nuclear extraction

Histones were isolated as previously described^19^. In summary, after nuclei isolation, histones were acid extracted with H_2_SO_4_ for three hours, following by precipitation with trichloroacetic acid (TCA) overnight. The resulting pellets were washed first with acetone containing 0.1% of hydrochloric acid (HCl) followed by a wash in acetone alone. Finally, samples were resuspended in ultrapure water and protein concentrations were determined by Bradford assay (BioRad). To generate nuclear extracts, ~1×10^6^ cells, were lysed in hypotonic buffer (10mM Tris pH 8, 2mM MgCl_2_, 24mM CaCl_2_, 0.3M sucrose, 0.5% NP-40 and protease inhibitors) for five minutes on ice. The nuclei were pelleted at 500g for 5 minutes and washed with PBS. Nuclear proteins were released on ice for 30 minutes in nuclear extraction buffer (10mM HEPES pH 7.9, 0.75mM MgCl_2_, 500mM NaCl, 0.1mM EDTA, 12.5% glycerol) supplemented with 2.5 U/ul benzonase (Millipore) and protease inhibitors (Bimake).

### Antibodies

All antibodies used in this study are listed in Supplementary Table 1.

### Co-immunoprecipitation (Co-IPs) and protein identification by mass spectrometry

Co-IPs were performed as previously described ^48^. Briefly, 2×10^7^ cells were lysed for 1hr at 4°C in IP buffer (20mM Tris pH 7.5, 137mM NaCl, 1mM CaCl_2_, 1% NP-40, 10% glycerol, 1mM MgCl2 and 12U/ml Benzonase) supplemented with protease inhibitors (Bimake). Antibodies (FLAG, BAF180, BAF170, IgG) were conjugated to 30μl protein G magnetic beads (Thermo) in blocking solution (0.5% BSA in PBS) for at least 2hrs at 4°C. Cell lysate (~600μg) was added to the conjugated beads and incubated overnight at 4°C. IP samples were washed three times with IP buffer and eluted in SDS loading buffer for immunoblots or processed for MS analysis by on-beads digestion (FL-H2A and cH2A IPs) as follows. Before trypsin digestion, beads were resuspended in 50mM NH4HCO3, and reduction with DTT, alkylation with iodoacetamide and digested with Trypsin (Promega). Resulting peptides were dried by SpeedVac centrifugation and store at −80°C. For MS analysis, samples were resuspended in MS buffer A (0.1% formic acid in MS grade water) and analyzed by LC-MS. Samples were loaded onto a 17-cm in house C_18_ column (Reprosil-Pur) using a Dionex LC (Thermo Scientific) coupled to a QE-HF orbitrap mass spectrometer (Thermo Scientific). Peptides were eluted over a 90-minute gradient from 5-35% solvent B (solvent A: 0.1%FA, solvent B: 80% acetonitrile, 0.1% FA), the flow rate was set to 300nl/min. Data was obtained in data dependent acquisition (DDA) mode. The MS1 was acquired over 300-1200 *m/z* with a resolution of 60,000, AGC target of 5e5, and maximum injection time of 100 ms. For the MS2, the top 25 most intense ions were selected for MS/MS by high-energy collision dissociation (HCD) at 27 NCE, with a resolution of 30,000, AGC target of 1e5, and maximum injection time of 150 ms. Raw files were processed using MaxQuant (v 1.6.0.16) ^49^ using *M. Musculus* database (Uniprot, April 2017). For MS/MS database search the precursor mass tolerance was set to 4.5ppm and the product mass tolerance to 0.5 Da. Two trypsin missed cleavages were allowed. Carbamidomethyl (C) was set as a static modification while oxidation (M) and acetylation (protein N-terminus) were selected as variable modifications. Proteins were quantified using label-free quantification (iBAQ). The protein false discovery rate (FDR) was filtered for < 0.01.

### Mononucleosme IP and histone PTMs analysis by mass spectrometry

Cells were lysed in Buffer A (10mM HEPES pH 7.9, 10mM KCL, 1.5mM MgCl_2_, 0.34M sucrose, 10% glycerol 1mM DTT, 0.1% Triton-X and protease inhibitors) on ice for 5min. After centrifugation, pellets were washed with Buffer A without Triton-X. Soluble nuclear proteins were released for 15min on ice in no salt buffer (3mM EDTA, 0.2mM EGTA, 0.1mM DTT). Mononucleosomes were obtained by digesting the chromatin in 30 units of Micrococcal nuclease (Mnase)(Roche) in digestion buffer (50mM HEPES, 2mM CaCl_2_ 0.2% NP-40) for 10min at 37 °C. Chromatin was briefly sonicated at 20% amplitude. Sample was further clarified by centrifugation and DNA was examined by agarose gel electrophoresis. Monucleosomes were dialyzed against Buffer D (20mM HEPES pH 7.9, 20% glycerol, 0.2mM EDTA 0.2% Triton X, and protease inhibitors) for 2hr at 4°C. Following dialysis, ~100ug of mononucleosomes were incubated with FLAG-M2 beads (20μl) overnight at 4°C. After three washes with Buffer D (Buffer A+100mM NACL), samples were eluted with 3X FLAG peptide (0.33mg/ml) on ice for 45min and then analyzed by immunoblotting and MS. For MS analysis, samples were run on a SDS gel and bands between 10-25kDa were excised. Histone peptides were derivatized with propionic anhydride described in Sidoli and Garcia 2017^50^. In summary, after two rounds of in gel derivatization, samples were digested with 12.5ng/μl of trypsin (Promega) overnight. Peptides were eluted form the gel after several iteration of hydration with 50mM NH4HCO3 and dehydration with 100% acetonitrile. Samples were dried by SpeedVac centrifugation and resuspended in 50mM NH4HCO3 for two more rounds of derivatization. Peptides were desalted and analyzed by LC-MS described above. Peptides were eluted over a 60-minute gradient from 5-35% solvent B (A:0.1% FA, B:80% acetonitrile, 0.1% FA), with a flow rate of 300nl/min. Data was obtained by data independent acquisition (DIA). The MS1 was done in high resolution (120, 000) from 300-1,*100m/z* w. For MS2, 16 DIA scans with *50m/z* windows were acquired using HCD (NCE =35). Raw files were processed using EpiProfile ^51^. These experiments were performed in triplicate.

### Peptide pulldowns and protein identification by MS

Histone peptide pulldown assays were done as previously described by Vermeulen ^52^ with minor modifications. In brief, nuclear extracts were prepared by lysing mESC in buffer A (10mM HEPES pH 7.9, 1.5mM MgCl_2_, 10mM KCl) supplemented with protease inhibitors. Cells were incubated on ice for 10min and pelleted at 400xg for 5min. Subsequently, cell pellets were further homogenized in Buffer A supplemented with 0.15% NP-40 and protease inhibitors. After nuclei were pelleted after spinning at 3200xg for 15min, nuclear proteins were released for 1hr at 4°C in Buffer C (420mM NaCl, 20mM HEPES pH 7.9, 20% glycerol, 2mM MgCl_2_, 0.1% NP-40 and protease inhibitors). Protein concentration was determined by Bradford assay.

Synthetic peptides were purchased from GenScript. Peptides were designed to include H2AK5ac, H2AK9ac, H2AK5acK9ac, H3K14ac and H4K5acK8ac, as well as corresponding control peptides without lysine acetylation. All peptides were biotinylated at the C terminus. The exact sequences of the peptides are listed in Supplementary Table 2. Each peptide (~25ug) was bound to 75μl of Dynabeads MyOne C1 (ThermoFisher) for 20 minutes at room temperature in peptide binding buffer (150 mM NaCl, 50 mM Tris pH 8, 0.15 %NP-40). Nuclear lysate was diluted to 0.6μg/ul in protein binding buffer (150 mM NaCl, 50 mM Tris pH 8, 0.15% NP-40, 0.5 mM DTT, 10 uM ZnCl_2_ and protease inhibitors). Nuclear lysates (~500ug) were incubated with conjugated peptides overnight at 4C. After 5 washes with wash buffer (protein binding + 350 mM NaCl) samples were eluted with SDS loading dye for MS or immunoblot analysis. For MS analysis, samples were resolved by SDS-PAGE (in 4-12%NuPage). Lanes were cut from 10-250kDa and divided into 4 different fractions. Before trypsin digestions, proteins were reduced with 5 mM DTT for 45min at 55°C and alkylate with 55 mM iodoacetamide at room temperature for 30min. Samples were digested overnight, and peptides were extracted as previously mentioned. Peptides were identified by MS as described in the Co-IPs section.

### H2A Top-down analysis by LC-MS

Prior to mass spectrometry analysis, individual histones were fractionated by reversed-phase liquid chromatography (RP-HPLC) as previously described ^53^. Briefly, ~200μg of purified histones were fractionated using Vydac C18 column (4.6 mm inner diameter, 250 mm length and 5 μM particle size). Samples were eluted using a 100min linear gradient (30-60%B). Solvent A consisted of 5% acetonitrile (ACN) in HPLC-grade water and .2% trifluoracetic acid (TFA), where Buffer B was 95% ACN and 0.2% TFA. Fractions corresponding to H2A.1 were collected based on known retention time and characteristic peaks (Supplementary Fig 1 A). Samples were then dried using a SpeedVac and stored at −80 °C. Fractions corresponding to H2A.1 were resuspended in 0.1% formic acid (FA) to a final concentration of 1μg/μl. Samples (2 μg) were loaded onto a 75-um (inner diameter) x 20 -cm C18 column (Reprosil-Pur Germany) for nano-LC. Proteoforms were eluted over 60min using a non-linear gradient (10-40B%). Solvent A consisted of 0.1% formic acid while solvent B was 80% acetonitrile (ACN). nLC was coupled to a hybrid Orbitrap Fusion Mass Spectrometer equipped with electron transfer dissociation (ETD) fragmentation (Themo). The acquisition was done in data dependent mode with a 3 second cycle time and using high resolution of MS1 (120,000 at 200=m/z) and tandem MS2 (30,000). For MS1 and MS2 the AGC target was set to 2.0E5 with a max injection time of 100 ms for MS1 and 300 ms for MS2. Charge states 5-21 were selected for MS2 fragmentation with ETD, finally 3 micro scans were averaged to yield high resolution MS/MS spectra. Data analysis was performed as described in Sidoli *et al.* with minor modifications. ^54^. In brief, after spectral deconvolution with Xtract (Thermo), files were search with Mascot (v2.5 Matrix Science). Acetylation (K) and methylation (KR) were included as dynamic modifications. No enzyme was selected for digestion. Results files were further process using IsoScale^54^ with filtering for unambiguous identification. All cleavage sites were normalized to the abundance of the full length H2A, manually extracted using Xcalibur (Thermo Fisher Scientific)

### Quantitative Proteomics with Multiple Reaction Monitoring (MRM) of H2A acetylation

For histone UHPLC-MRM-MS analysis, 20μg of histone was derivatized as previously described^41^. After two rounds of derivatization, samples were digested with trypsin overnight. Tryptic peptides were derivatized twice more and peptides were desalted by stage tipping ^41^. The multiple reaction monitoring (MRM) method was developed on a Vanquish liquid chromatography system and TSQ Altis triple quadrupole mass spectrometer (Thermo Scientific) to quantify H2A tail acetylation. Using 25 pmol of heavy labeled AQUA standard peptides, targets were scheduled with using a 15min gradient, peptides were loaded onto an Accucore 1.5μm x 15cm C_18_ column (Thermo Scientific) heated to 60°C at a flow rate of 300 μ<u>l</u>/min. Using an aqueous solution of 0.1% formic acid as Buffer A an organic solution of 80% acetonitrile with 0.1% formic acid as Buffer B, the gradient began at 5% Buffer B and increased to 50% Buffer B by 13.9min, then increased to 95% Buffer B at 15.0min where it remained for 2 minutes, and finally decreased to 5% Buffer A at 17.1min where it remained for 6 minutes. Peptides were introduced into the mass spectrometer by H-ESI source in positive mode at 3700V, sheath gas of 25Arb, auxiliary gas of 4Arb, sweep gas of 1Arb, ion transfer tube temperature of 325°C, and vaporizer temperature of 350°C. A final method was created using the previous parameters coupled to multiple reaction monitoring with a total number of 164 transitions, dwell time of 2.5ms, Q1 resolution of 0.4 Da, Q3 resolution of 0.7 Da, and at least ten points across each peak were quantified. Calculation of the heavy to light ratio of each peptide was carried out in Skyline ^55^. All experiments were done in triplicate.

### Chromatin Immunoprecipitation followed by sequencing (ChIP-seq)

ChIP-seq was done as previously reported ^48^. Briefly, 1×10^7^ mESCs were fixed in 1% formaldehyde in PBS for 10 minutes and then quenched with 125 mM glycine for 5min. Chromatin was sonicated with a S220 Focused-ultrasonicator (Covaris) for 15min and immunoprecipitated with anti-H2AK9ac (Abcam). Antibodies were conjugated to protein G Dynabeads (Thermo). After washing and elution, samples were incubated overnight at 65°C to reverse cross-links. DNA was purified after treatment with RNAse (Thermo) and Proteinase K (Thermo). Purified DNA was quantified using Qubit dsDNA kit (Thermo) and 50ng was used to prepare sequencing libraries. Input samples were also included and prepared using the same protocol. NEBNext Ultra DNA library kit for Illumina (New England Biolabs) was used to prepare the libraries and the quality was assessed by Agilent BioAnalyzer 2100 (Agilent). Quantification was done using KAPA library quantification kits (KAPA Biosystems) or NEBNext library Quant kit (New England Biolabs). For each condition, two independent replicates were included. Single-end sequencing (75bp) was performed on a NextSeq 500 platform. (Illumina). ARID2 and PBRM1 ChIP-Seq was done as reported by Michel et. al^56^.

Briefly, 2×10_7_ mESCs were fixed with 1% formaldehyde for five minutes and quenched for an additional five minutes. After nuclear extraction, chromatin was sonicated with a S220 Focused-ultrasonicator (Covaris) for 12 min and immunoprecipitated with anti-PBRM1 (Bethyl) and anti-ARID2 (Bethyl). After washing and elution, samples were incubated overnight at 65°C to reverse cross-links. DNA was purified after treatment with RNAse (Thermo) and Proteinase K (Thermo). Using the Qiagen minElute PCR purification kit (Qiagen). Purified DNA was quantified using Qubit dsDNA kit (Thermo) libraries using the KAPA Hyper Prep kit Sequencing was performed at the Genome Sequencing Facility in St. Jude Children’s Research Hospital on Illumina HiSeq platform in single-end mode with 50 bp per read. Reads were checked for sequencing quality with FastQC (v0.11.2). After that, reads were aligned to the mouse reference genome (mm9) using Bowtie2 (v2.2.9) with soft-clipping allowed. SAMtools (v0.1.19) was used to filter out non-primary alignments, alignments with mapping quality score less than 10, PCR duplicates, and alignments on non-chromosome contigs or ENCODE blacklist regions. Peaks were detected for each sample against the corresponding input control using SICER (v1.1) with default parameters. Bedtools (v2.27.1) was used to generate bedgraph files for visualization on genome browser, in which each sample was normalized to 10 million reads per library with corresponding input subtracted.

### Whole transcriptome analysis by RNA-seq

RNA was extracted utilizing RNAeasy Kit (Qiagen) following manufacturer’s instructions. Treatment with DNase for 15 minutes was included to degrade all genomic DNA. Libraries were prepared using NEBNext Poly(A) mRNA magnetic isolation module and NEB Ultra Directional RNA Library kit for Illumina (New England Biolabs). Library quality, quantification and, sequencing was done as described above. RNA-seq reads were checked for sequencing quality with FastQC (v0.11.2). After sequencing, reads were aligned to the mouse reference genome (mm9) using STAR (v.2.3.0e) with default parameters. SAMtools (v0.1.19) was used to filter out alignments with mapping quality score less than 10 and alignments on mitochondria or non-chromosome contigs. Finally, FeatureCounts (v.1.6.2) was used to generate a matrix of mapped fragments per RefSeq annotated gene. Differential gene expression analysis was performed using the DESeq2 R package (v.1.16.1) with an FDR□cutoff of□0.05 and log2 fold change cutoff of 1.5X. Bedtools (v2.27.1) was used to generate bedgraph files for visualization on genome browser, in which each sample was normalized to 10 million reads per library.

### GO analysis

All gene ontologies analysis were done using GENEONTOLOGY at http://geneontology.org/

## Data availability

Mass spectrometry raw files are deposited in Chorus repository project number 1698. Additionally, all ChIP and RNA seq files are in the NCBI Gene Expression Omnibus under the following accession number GSE162896.

## Acknowledgments

The authors thank Dr. Shelley Berger for providing plasmids used here. We also thank Dr. Kate Alexander and Dr. Enrique Lin Shiao for their scientific input and discussions. we acknowledge the following grants NIH GM110174, CA196539 and NS111997. SS gratefully acknowledges the Leukemia Research Foundation (Hollis Brownstein New Investigator Research Grant), AFAR (Sagol Network GerOmics award), Deerfield (Xseed award), Relay Therapeutics, Merck and the NIH Office of the Director (1S10OD030286-01). Additionally, BAG acknowledges funding from a St. Jude Children’s Hospital Consortium.

## Author contributions

B.G. and M.C. conceived the idea for this project and designed experiments. M.C. wrote the manuscript with essential contribution from EGP. M.C also designed experiments and performed experiments presented here. J.C assisted and designed quantitative QQQ method to analyzed H2A acetylation. Y.L performed computation analysis for sequencing experiments with significant contributions from PJL. Z.Z and C.R assisted and designed sequencing experiments with invaluable contributions from Y.P. C.L, and S.S assisted with Top-Down analysis and LC-MS method development. All authors discussed the results and the manuscript.

## Supplementary figure legends

**Supplementary figure 1. A** Representative chromatogram of reverse-phase fractionation (RP-HPLC) of acid-extracted histones from mESCs. Top-down quantification of proteolytic events found at day 0 **B**, 2-**C** and 4-**D.** Error bars represent standard deviation of 3 experiments. **E** Sequence alignment of H2A variants, yellow box shows sequence homology on the most abundant cleavage site of H2A after 4 days of differentiation.

**Supplementary figure 2**. Top-down quantification of proteolytic events in shSC (control) and shCTSL mESCs in before **A** and after 2 days of differentiation **B.** Only peptides found in 2 out of 3 replicates are reported. Bottom-up quantification of H2AK5ac **C**, H2K9ac **D** and H2AK5acK9ac **E** during ES differentiation, y axis shows the average of three biological replicates, error bars represent standard deviation. P-values were determined using unpaired Student t-test *p < 0.05, **p < 0.01, ***p < 0.001

**Supplementary figure 3. A** MS quantification of H2AK9ac in undifferentiated shSC control (blue) or shCTSL cells (orange) using synthetic peptides as standards. Error bars represent standard deviation of 3 experiments. **B** Distribution of H2AK9ac peaks across genomic features in undifferentiated cells shSC control (blue) and shCTSL cells (orange). ChIP seq of undifferentiated cells performed once). **C** Gene ontology of differentially downregulated genes from day 2 to day 4 of EBs. **D** Genomic distribution of H2AK9ac on day 4 of EBs formation in control shSC or after shCTSL cells.

**Supplementary figure 4. A** Peptide pull-down using unmodified H2A, H2AK5ac, H2AK9ac or H2K5acK9ac peptides. Pull-downs were carried out using nuclear extracts of undifferentiated ESCs, western blots were done against proteins that showed >2-fold enrichment by MS. **B** *in-vitro* peptide pull-down using recombinant BRD4 or recombinant BRD7. **C** Peptide pull-down of unmodified H4 and H4K5acK8ac, western blot against BRG1 and BRD4, well-known reader of acetylated H4. **D** Peptide pull-down using unmodified H2A, H2K5ac, K9ac, H2K5acK9ac, H3, H3K14ac, H4 and H4K5acK8ac peptides. Western blots were done against Oct4 (negative control) and TBP. **E** Venn diagram showing overlapping peaks between H2AK9ac and Arid2 of Pbrm1 **F.**

**Supplementary figure 5.** Gene ontology of proteins found to interact with cH2A **A** or FL-H2A **B** with FC > 0.5 and p-value > 0.05 using unpaired t-tests. Top 10 categories are shown. **C** Biological replicate of Co-IPs in Figure 5D.

**Supplementary figure 6. A** Simplified schematic of monoucleosme IP procedure. **B** Biochemical fractionation showing cH2A is chromatin associated. **C** Biological replicates of protein stability experiments described in figure 6 C and D.

**Supplementary table 1.** Antibodies used in this study

| Target | Source | Concentration/Application |
| --- | --- | --- |
| FLAG | Sigma F3165 | 3ug (Ip), 1:5000 WB |
| BAF180 | Millipore ABE70 | 5ug (Ip), 1:1000 WB |
| BAF180 | Bethyl A700-019 | 5ug ChIP |
| BAF170 | Active motif 61471 (IP) | 5ug (Ip) |
| BAF170 | Cell signaling (WB) | 1:1000 WB |
| ARID2 | Thermo Invitrogen pa5-5128 | 1:1000 WB |
| ARID2 | Bethyl A302-230A (ChIP-Seq) | 5ug ChIP |
| BRG1 | Abcam 110641 | 1:1000 WB |
| BRD7 | Cell signaling 15125s | 1:1000 WB |
| TBP | Abcam 133239 | 1:1000 WB |
| OCT4 | Abcam 19857 | 1:1000 WB |
| H2AK9ac | Abcam 177312 | 3ug ChIP |
| H2A-acidic patch | Millipore 07-146 | 1:1000 WB |
| Cathepsin L | R&D biosystems AF1515 | 1:1000 WB |
| GAPDH | Cell signaling | 1:10000 WB |
| BRD4 | Abcam 128874 | 1:1000 WB |
| GST | Cell signaling 2622s | 1:1000 WB |
| H3 | Abcam 1791 | 1:5000 WB |
| H2B | Cell Signaling | 1:1000 WB |
| IgG | Abcam 171870 | 5ug (Ip) |

**Supplementary table 2.** Synthetic peptides used in this study (20aa)

| Histone | Peptide sequence | Name |
| --- | --- | --- |
| H2A | Ac-SGRGKQGGKARAKA{Lys(Biotin)}TRSSR | H2A unmodified |
|  | Ac-SGRG{Lys-Ac}QGGKARAKA{Lys(Biotin)}TRSSR | H2AK5ac |
|  | Ac-SGRGKQGG{Lys-Ac}ARAKA{Lys(Biotin)}TRSSR | H2AK9ac |
|  | Ac-SGRG{Lys-Ac}QGG{Lys-Ac}ARAKA{Lys(Biotin)}TRSSR | H2AK5acK9ac |
| H4 | Ac-SGRGKGGKGLGKGGAKRHR{Lys(Biotin)} | H4 unmodified |
|  | Ac-SGRG{Lys-Ac}GG{LysAc}GLGKGGAKRHR{Lys(Biotin)} | H4K5acK8ac |
| H3 | ARTKQTARKSTGGKAPR{Lys(Biotin)}QL | H3 unmodified |
|  | ARTKQTARKSTGG{Lys-Ac}APR{Lys(Biotin)}QL | H3K14ac |

