## Supplementary figures for "Cleavage of histone H2A during embryonic stem cell differentiation destabilizes nucleosomes to counteract gene activation"

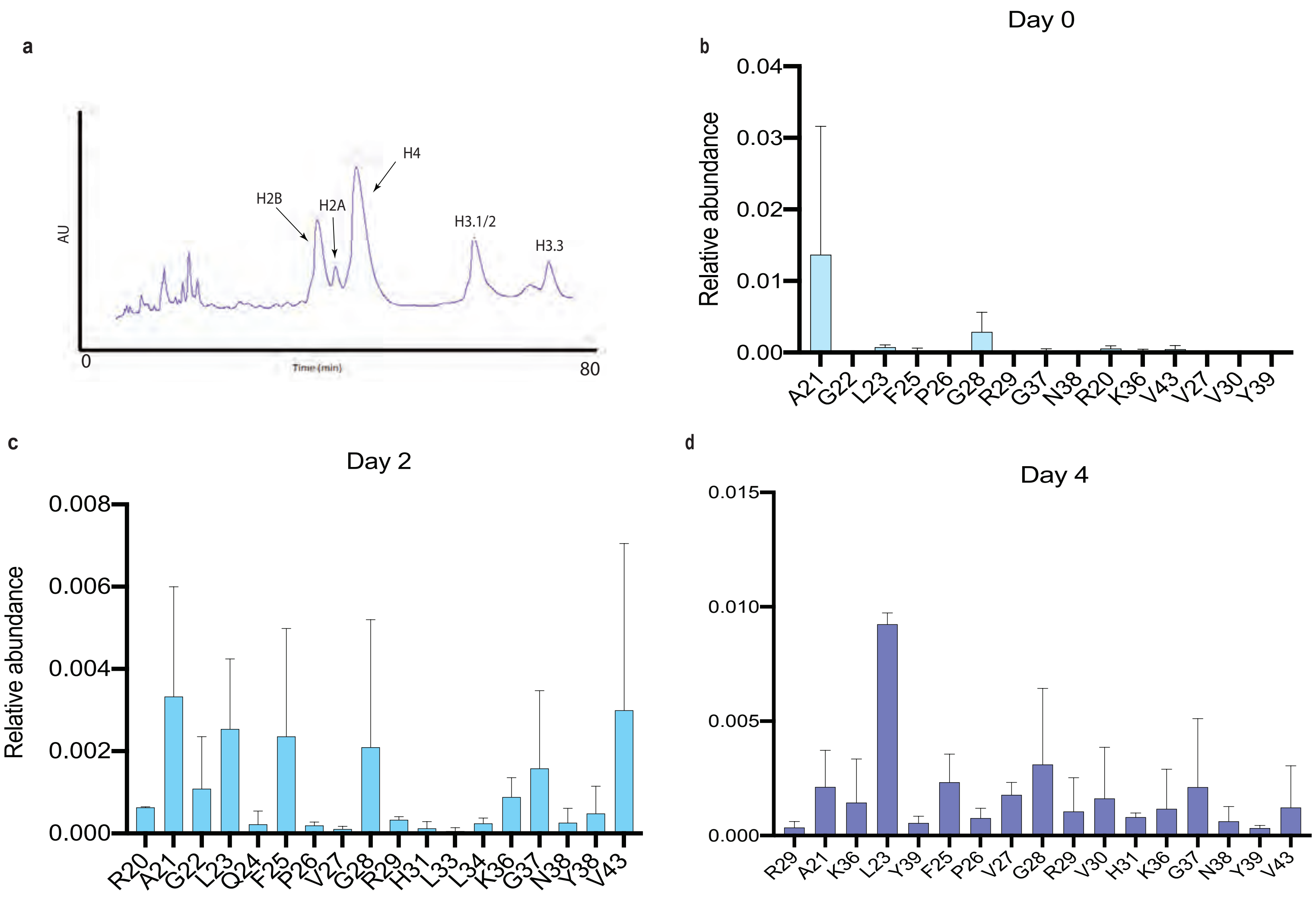


**e**

```
H2A2A_MOUSE MSGRGKQGGKARAKAKSRSSRAGLQFPVGRVHRLLRKGN-YAERVGAGAPVYMAAVLE
H2A1H_MOUSE MSGRGKQGGKARAKAKTRSSRAGLQFPVGRVHRLLRKGN-YSERVGAGAPVYLAAVLE
H2AX_MOUSE  MSGRGKTGGKARAKAKSRSSRAGLQFPVGRVHRLLRKGH-YAERVGAGAPVYLAAVLE
H2A3_MOUSE  MSGRGKQGGKARAKAKSRSSRAGLQFPVGRVHRLLRKGN-YSERVGAGAPVYLAAVLE
H2AB1_MOUSE ---     MARKRQRRRRRKVTRSQRAELQFPVSRVDRFLREGN-YSRRLSSSAPVFLAGVLE
H2A1K_MOUSE MSGRGKQGGKARAKAKTRSSRAGLQFPVGRVHRLLRKGN-YSERVGAGAPVYLAAVLE
H2AY_MOUSE  ---MSSRGGKKKSTKTSRSAKAGVIFPVGRMLRYIKKGH-PKYRIGVGAPVYMAAVLEYLTAE
H2AZ_MOUSE      GGKAGKDSGKAKTKAVSRSQRAGLQFPVGRIHRHLKSRTTSHGRVGATAAVYSAAILE
H2A1C_MOUSE MSGRGKQGGKARAKAKTRSSRAGLQFPVGRVHRLLRKGN-YSERVGAGAPVYLAAVLE
H2A1O_MOUSE MSGRGKQGGKARAKAKTRSSRAGLQFPVGRVHRLLRKGN-YSERVGAGAPVYLAAVLE
```

Supplementary figure 2

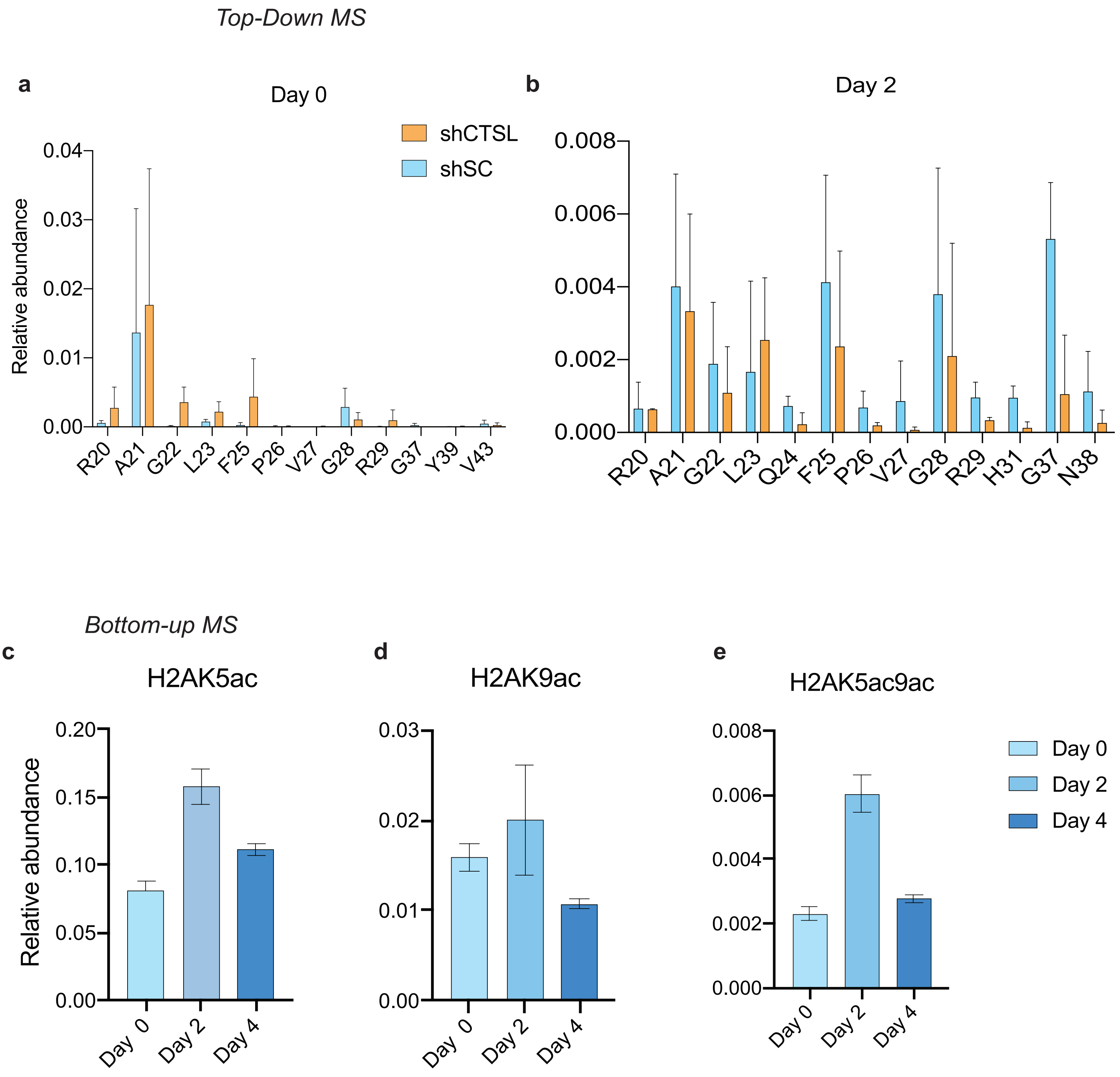

### Supplementary figure 3

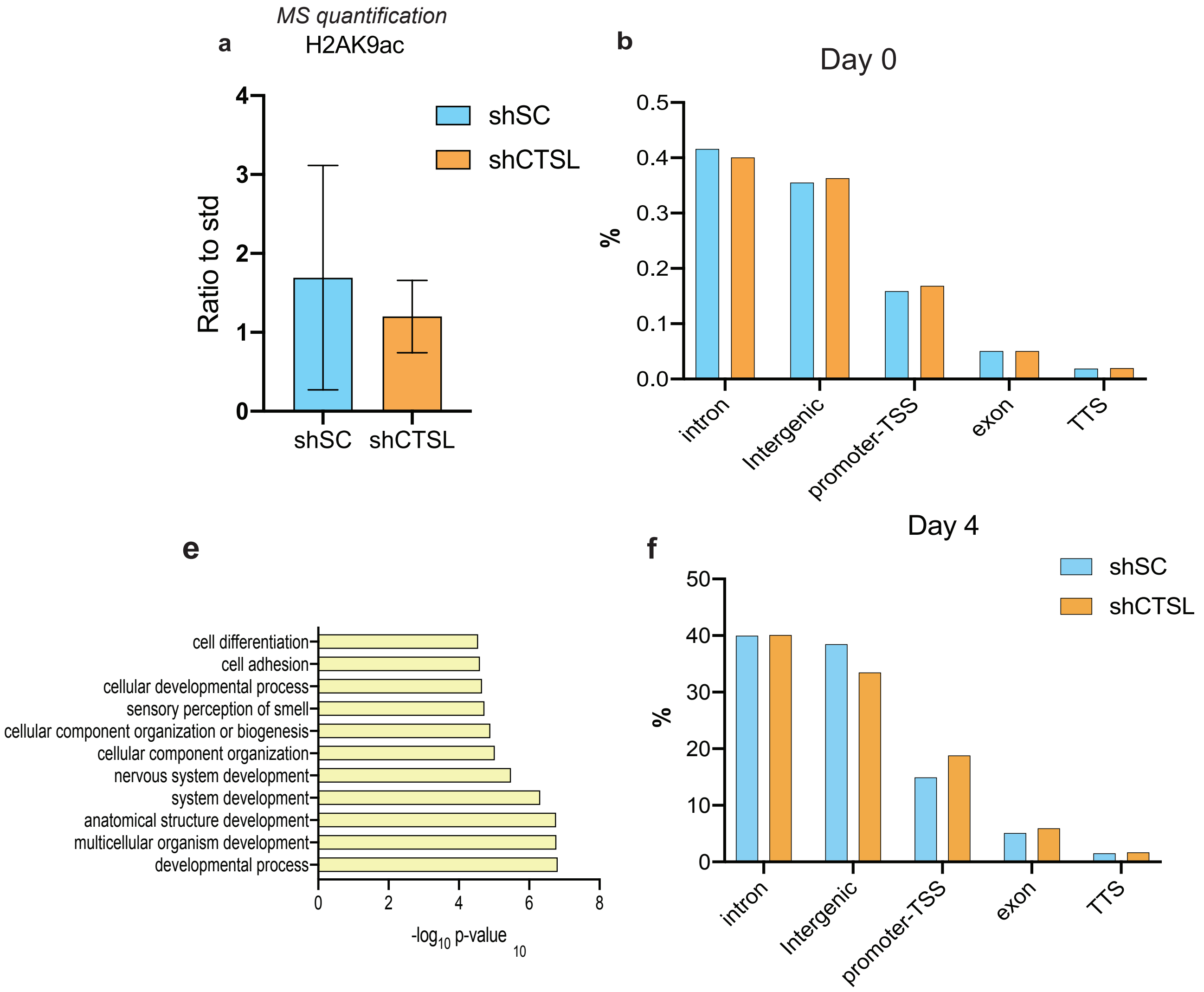

Supplementary figure 4

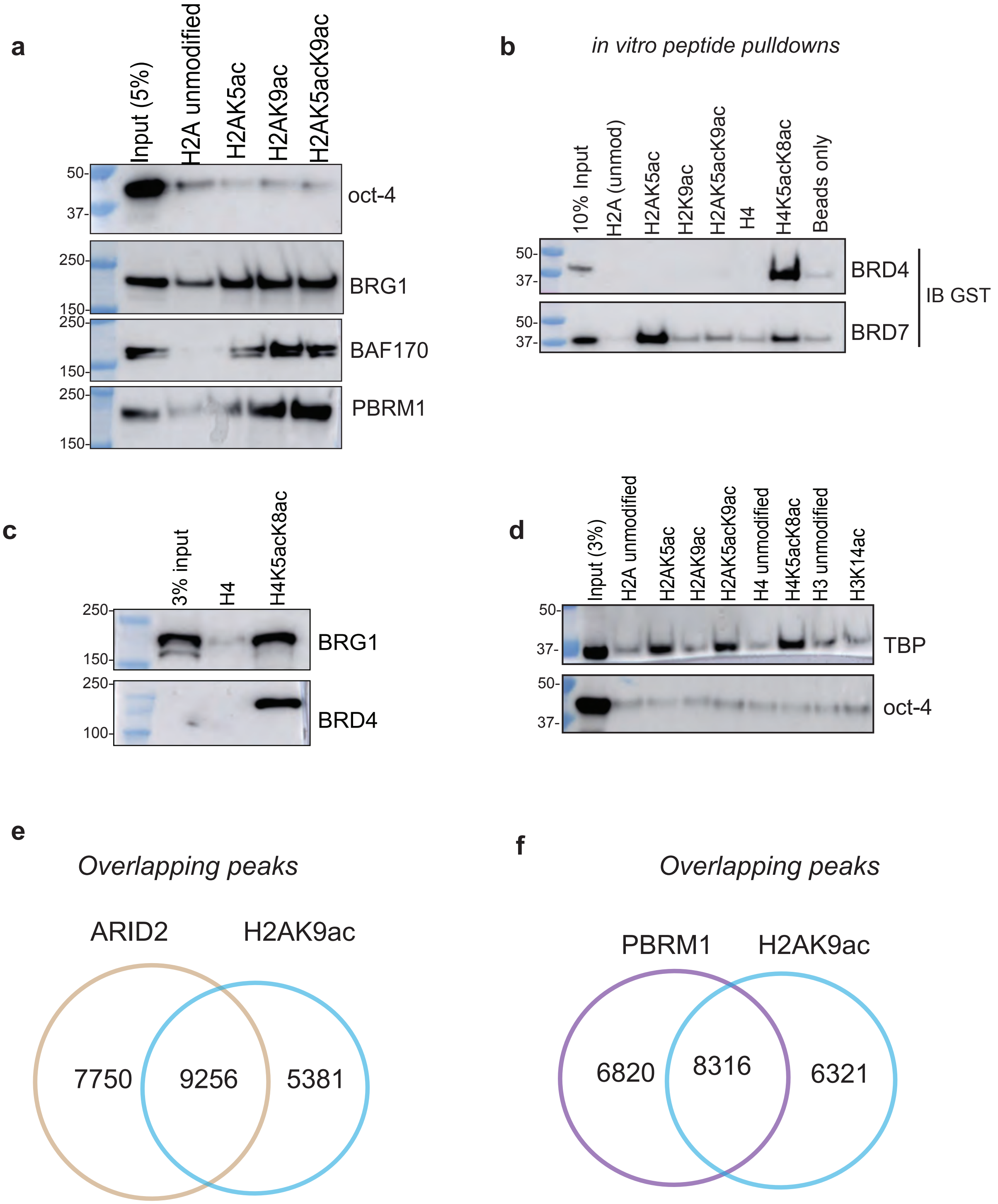

Supplementary figure 5

**a**

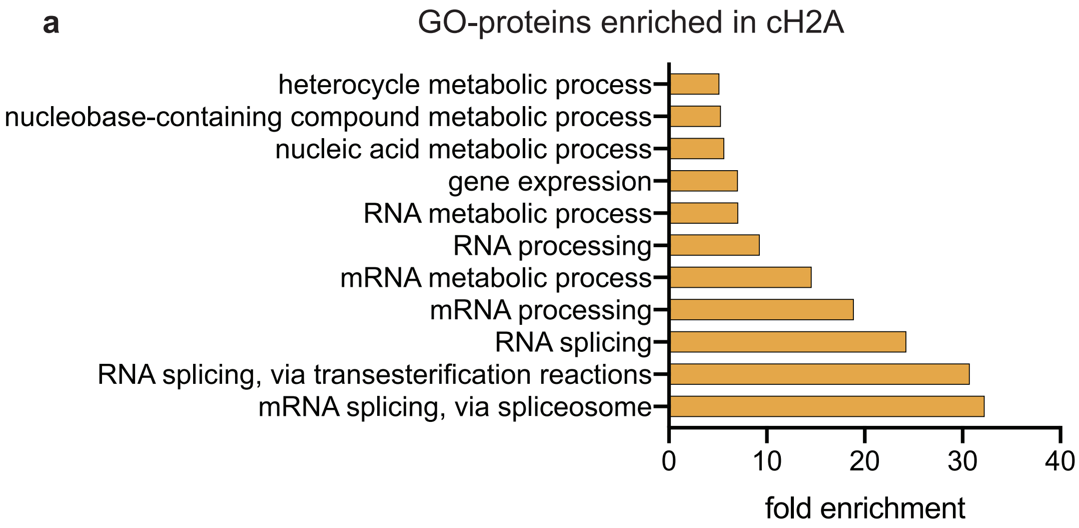


**b**

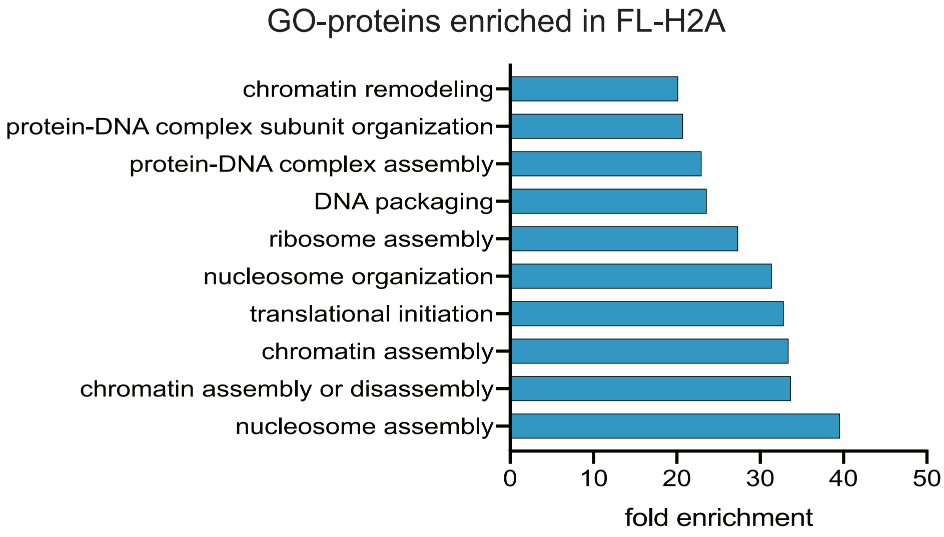


**c**

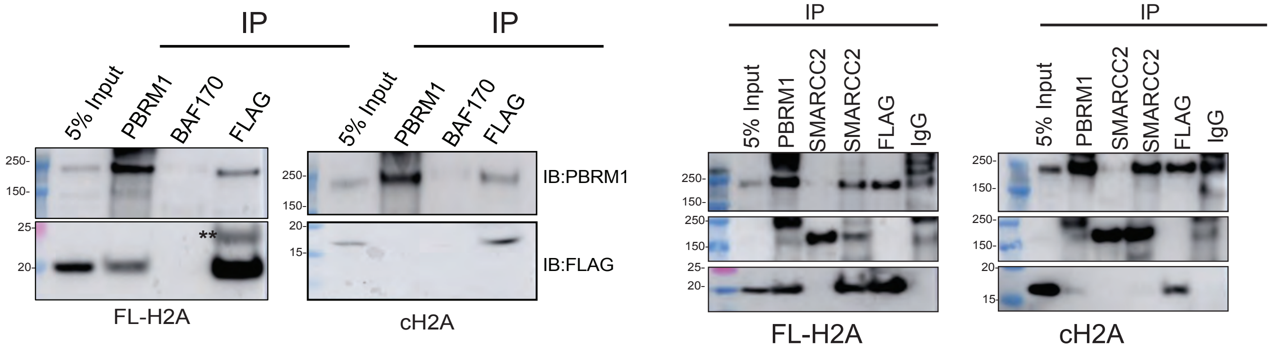

Supplementary figure 6

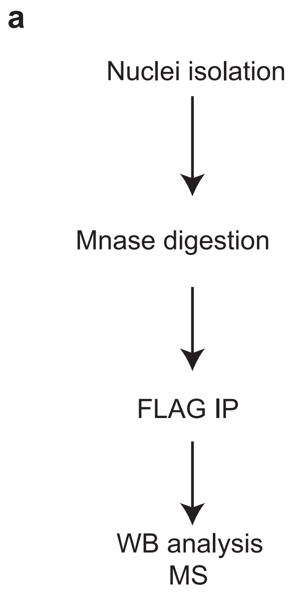


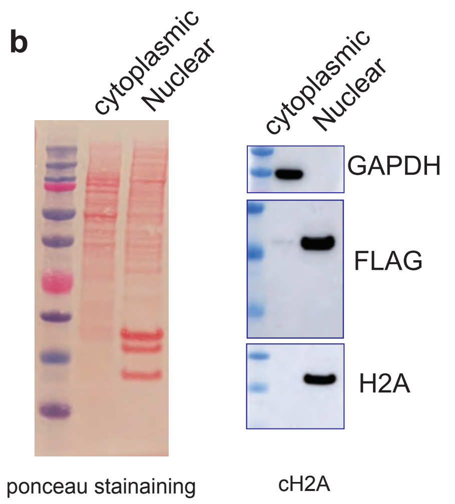


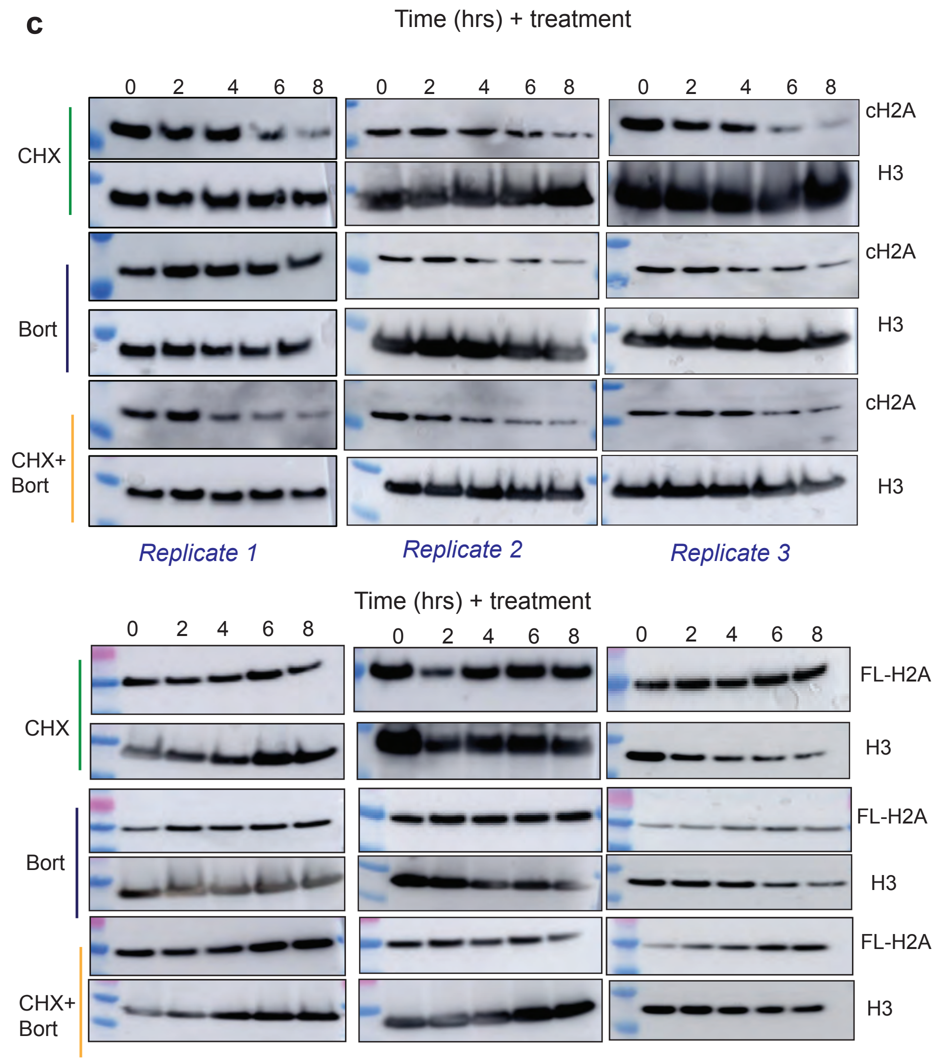
